# The Human Contraceptive Antibody Targets A Semenogelin Galectin-3 *N-*Linked Glycan Complex on the Sperm Surface

**DOI:** 10.64898/2026.10.13.757762

**Authors:** Sarah Dohadwala, Chaoshuang Xia, Shane Finn, Purna Shah, Shang-Chuen Wu, Hau-Ming Jan, Joseph A Politch, Kevin J Whaley, Sean R Stowell, Stacy Malaker, Catherine E Costello, Deborah J Anderson

**Affiliations:** Dept of Virology, Immunology, and Microbiology, Boston University, Boston, MA, United States of America; Center for Biomedical Mass Spectrometry, Boston University, Boston, MA, United States of America; Department of Chemistry, Yale University, New Haven, CT, United States of America; Department of Biology, Boston University, Boston, MA, United States of America; Joint Program in Transfusion Medicine, Brigham and Women’s Hospital; Department of Medicine, Boston University Chobanian & Avedisian School of Medicine, Boston, MA, United States of America; ZabBio Inc. San Diego, CA, United States of America

**Keywords:** sperm, galectin, contraception, antibody, CD52, semenogelin

## Abstract

The Human Contraception Antibody (HCA), is a monoclonal antibody that agglutinates and immobilizes sperm, and is the active component in a contraceptive product undergoing clinical trials. To date, the HCA epitope has been described as an *N*-linked polylactosamine glycan on CD52, but this assignment has been inferred by colocalization with anti-CD52 and S19 antibodies in western blots. Here, we report that HCA binds semenogelin-galectin-3 complexes associated with sperm. Semenogelins are gel-forming, highly glycosylated proteins that are cleaved into smaller segments that adhere to sperm during semen liquefaction. We isolated HCA binding partners from seminal plasma and sperm, and semenogelin was identified in the binding fraction by mass spectrometry. Western blots demonstrated that HCA binds to a large smear of seminal plasma proteins with a higher apparent molecular mass than CD52. Flow cytometric analysis of deglycosylated sperm demonstrated that HCA binding requires sperm surface *N*-linked glycans. Despite this, HCA failed to bind any glycans represented on a comprehensive microarray. In addition to semenogelin, Galectin-3 was identified in the HCA purified sperm lysates, supporting a model in which HCA’s binding site contains semenogelin-derived glycopeptides tethered to sperm via galectins bound to an *N-*linked glycan. We found that *O*-linked glycans moieties on semenogelin contain LacNAc motifs that are recognized by galectins. Finally, depletion of sperm surface galectins reduced HCA-mediated agglutination, and this was rescued by addition of recombinant galectin-3/semenogelin-2 complexes. Collectively, our findings suggest that HCA targets an antigen that contains a semenogelin-galectin-3 complex anchored to the sperm surface via an *N*-linked glycan.

**Summary sentence:** We provide evidence that the Human Contraception Antibody binds to a semenogelin derived glycopeptide complex that attaches to the surface of sperm via a galectin mediated interaction; it remains unclear whether CD52 is part of the complex.

Graphical Abstract

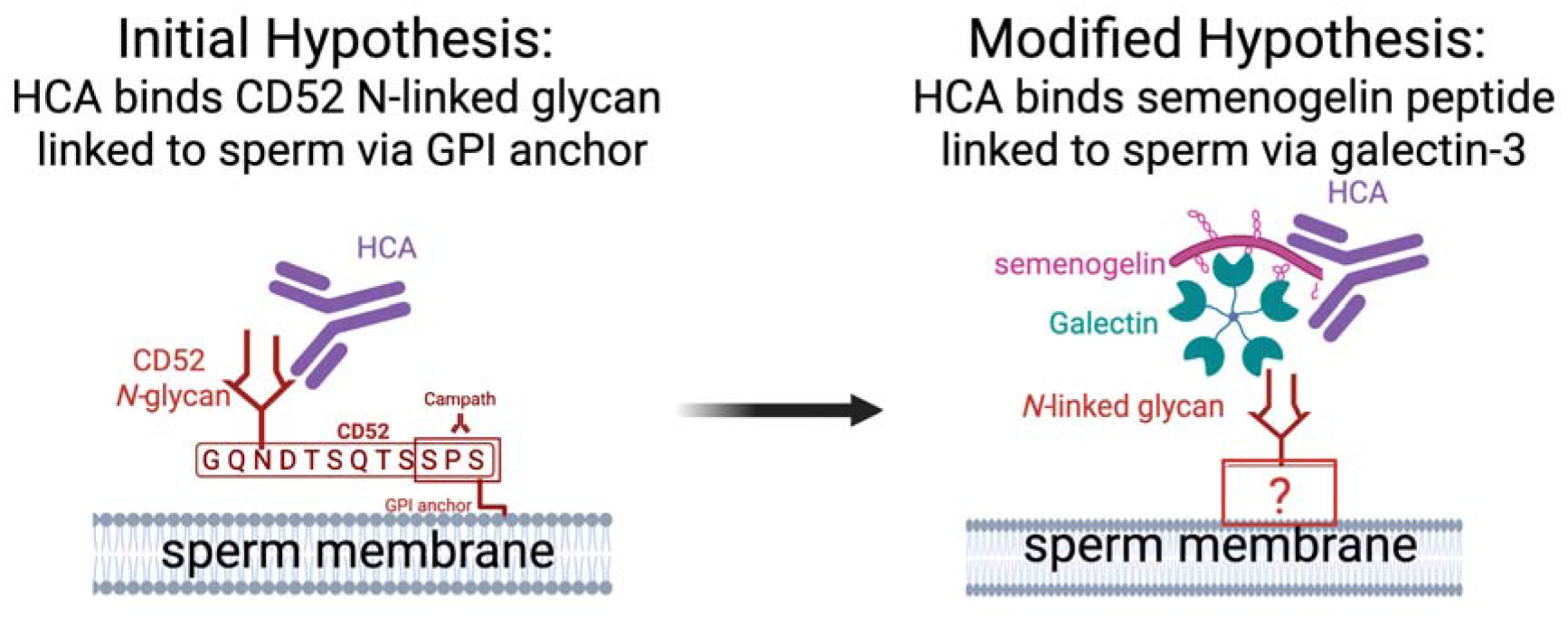

## Introduction

The Human Contraception Antibody (HCA) is a monoclonal antibody that is being developed as a non-hormonal contraception candidate (*1*). Several interesting properties of HCA made it an ideal candidate to develop as a contraceptive. Firstly, the antibody potently agglutinates and immobilizes sperm, preventing their movement into the cervix and thus their ability to access the uterus(*1*). Secondly, HCA appears to bind a male reproductive tract (MRT)-restricted epitope present in semen and particular regions of the MRT, including the cauda epididymis, and does not bind to vaginal cells (*1*). Because HCA is a human antibody, it does not induce toxicity or inflammation in the vaginal tract, a shortcoming of several other topical contraception methods (*1*, *2*).

To develop HCA as a monoclonal antibody drug, the sequence of the antibody was engineered from the original IgM sequence of H3C4, an antibody originally cloned from the blood of an infertile woman by Isojima et al (5). The antibody Fab sequence was inserted into a human IgG1 backbone, and produced at GMP grade in Nicotiana plants, which have a defined, homogenous Fc glycosylation profile (*3*). This high-quality antibody was formulated into a polyvinyl alcohol film for topical delivery to the vaginal tract, and its safety and surrogate efficacy were assessed in a Phase I clinical trial (2). Excitingly, the film containing HCA was safe, and the surrogate efficacy measure, a post-coital test, indicated that the women enrolled in the study (N = 8) had virtually no motile sperm in their cervical mucus when intercourse occurred after film insertion (2). The antibody is now entering additional Phase I and Phase II trials for further study of its safety, pharmacokinetics and dose finding.

It is exciting that the antibody works well *in vivo*, and with a relatively unique mechanism. While it is known that the contraceptive functions of HCA are reliant on agglutination and complement mediated immobilization, exactly how HCA mediates these functions is unknown (*4*). To date, the binding epitope of HCA has been described as glycan dependent (1, 5). The glycan epitope of HCA was originally defined as an internal repeat polylactosamine motif by thin layer chromatography of glycolipid standards derived from placenta, though important details of the methodology, such as which concentrations of HCA were used or detailed characterization or purity of the placenta-derived glycolipid standards are not described (5). These limited data suggest that the HCA binding site on the sperm surface may also include human glycolipids in addition to glycoproteins. In general, antibodies that interact with glycans are relatively understudied due to limitations in methods for synthesizing specific glycan antigens chemically (*6–8*). Follow up studies proposed the *N-*linked glycan of CD52 (referred to in the literature as SAGA-1) as a potential HCA epitope based on microsequencing and cross-immunoblotting of peptides affinity purified by S19, a mouse monoclonal antibody believed to share an epitope with HCA (9). To date, the literature continues to define the protein epitope of HCA by colocalization on western blots with a CD52 binding antibody, Campath (alemtuzumab) (1, 9). CD52 is a small GPI-anchored peptide on the surface of sperm that contains highly heterogeneous *N*-linked and *O*-linked glycan moieties (9). CD52 also interacts with several proteins on the surface of sperm that are important in reproductive processes, including eppin and semenogelin (10, 11, 41). Colocalization on blots cannot definitively assign HCA, S19, and CAMPATH the same antigen, given that several different proteins can be identified from the same apparent molecular weight on gels and western blots.

Given incomplete characterizations of the HCA epitope in the literature, we sought to define the glycoprotein binding partner of HCA in greater detail using modern mass spectrometry and classical biochemistry methods. To date, the sequence of HCA has not been successfully modified for substantial increased affinity to its antigen. While multimerization of its Fab regions has been pursued as a strategy to increase its potency, much potential remains in using techniques such as yeast surface display to develop more potent second-generation HCAs that can agglutinate and immobilize sperm at lower antibody concentrations. The limited characterization of the HCA target antigen is a clear roadblock in developing second generation therapeutics because all high throughput antibody affinity maturation methods require the production of pure antigen for use in assays. Understanding the binding site of HCA in greater depth will enable tool development to assess HCAs in biologically complex clinical material, develop second generation contraceptive antibodies, and understand the biological mechanism of sperm agglutination and immobilization. For these reasons, we repeated several key biochemical experiments from the literature and pursued a thorough investigation of the HCA binding site. Our investigation spans several orthologous methods, using immunoprecipitation followed by mass spectrometry, western blotting, flow cytometry, competition-agglutination experiments, glycan binding arrays, mass spectrometry-based glycopeptidomics, and galectin depletion kinetic agglutination followed by recombinant rescue, to determine the HCA binding site with higher confidence.

## Materials and Methods

### Materials

#### Collection and processing of human sperm

Semen was acquired from healthy donors with the approval of BUMC Institutional Review Board (IRB) under protocol #H36843. Samples were liquefied for 20 minutes at 37 °C, then motile sperm were separated from whole semen on a 90% ISolate density gradient (FUJIFILM Irvine Scientific; Santa Ana, CA, USA) by centrifugation at 300g for 20 minutes. The motile sperm pellet was resuspended in Multipurpose Handling Medium (MHM; FUJIFILM Irvine Scientific; Santa Ana, CA, USA). Sperm concentration and motility of both whole semen and isolated motile sperm were assessed using a Computer-Assisted Sperm Analysis system (CASA; Human Motility II software, CEROS II, Hamilton Thorne, Beverly, MA, USA). Samples with low concentrations (<10 million/mL) or motility (<40%) post-processing were not used.

#### Preparation of sperm lysates

Sperm lysates were prepared by pelleting motile sperm, resuspending in RIPA lysis buffer, and shearing through a narrow diameter syringe.

#### Preparation of sperm lysates lacking *N*-linked glycans

Intact sperm were treated with PNGase F (New England Biosciences) or PI-PLC (Invitrogen Cat # P6466) for 1 hour at 37 °C at a concentration of 35 million sperm per mL, in reaction volumes of 40 μL. Lysates were incubated with PNGase F overnight at 37 °C as per New England biosciences’ protocol.

#### Preparation of seminal plasma

Unliquefied seminal plasma (USP) was prepared by immediately diluting whole semen 1:5 in ice cold 4M urea, RIPA lysis buffer and 1mM PMSF to inhibit endogenous protease activity. Samples were centrifuged to pellet cell debris, and the supernatant was kept as USP. From processed semen samples, liquefied seminal plasma (LSP) was prepared by collecting the plasma fraction after centrifugation that remains above the Isolate density gradient. This fraction was then centrifuged to remove cell debris, and the supernatant was kept as LSP.

#### Recombinant semenogelin

Recombinant semenogelin-1 produced in HEK293 cells (Abcam, Cat# ab153783) and semenogelin-2 produced in mammalian cells (MyBioSource, Cat#: MBS963301) were purchased and reconstituted as per manufacturer protocols.

#### Recombinant Galectin-3

Plasmids encoding human galectin-3 were transformed into *E. coli* BL21 (DE3) and expressed as previously described (39, 40). Briefly, transformed bacteria were cultured in LB broth containing 100 μg/mL ampicillin at 37 °C. At mid-log phase, protein expression was induced by 1 mM isopropyl β-D-1-thiogalactopyranoside (IPTG). After 20 h of induction at 16 °C, bacterial culture was harvested by centrifugation and resuspended in bacterial lysis buffer consisting of PBS with 14 mM 2-mercaptoethanol (2-ME), 60 μl ribonuclease A, 60 μl DNase I, 60 μl lysozyme, and two protease inhibitor cocktail tablets (Roche). The suspension was passed through a cell disruptor, and the lysate was centrifuged at 17,000 rpm at 4 °C for 1 h twice. The supernatant was applied to a lactosyl-Sepharose affinity chromatography column at 4 °C. For elution, PBS containing 14 mM 2-ME and 100 mM lactose was used.

#### Antibodies

Human Contraception Antibody (HCA) was produced by ZabBio (San Diego, CA, USA) and KBio (Owensboro, KY) as previously described (1). HCA is a human IgG1 produced in glycoengineered *Nicotiana benthamiana*. Semenogelin-1 (ab139405) and Semenogelin-2 (ab244821) monoclonal antibodies were purchased from Abcam. Human IgG1 CAMPATH (Cat #: LT200-1MG) was purchased from Leinco, Mouse IgG1 CAMPATH (Cat #: PIMA547806) was purchased from Invitrogen, and rat IgG1 CAMPATH (Cat# MA5-16999) from ThermoFisher Scientific. Galectin-3 antibody was purchased from Abcam (ab76245).

### Methods

#### Sperm flow cytometry

Sperm were incubated with PNGase F (NEB), PI-PLC (NEB), or media alone for 2 hrs and then were fixed in formaldehyde. Fixed sperm were stained with 5 μg/mL HCA followed by goat anti-human secondary antibody (1:500 dilution) for 1 hour. Sperm were washed thrice between steps in PBS containing 1% BSA and 2mM EDTA. Fixed sperm were analyzed using a Cytek Aurora full spectral flow cytometer. Cells were gated by forward and side scatter and doublets were removed by gating the linear FSC-A vs FSC-H in FlowJo software.

#### Sperm kinetic agglutination assay

A kinetic agglutination assay was performed as previously described (37). A 2 µL sperm sample at a concentration of 35 million per mL was spotted then visualized at 10x magnification. Then, 2 µL of media containing HCA was mixed into the spot, and agglutination was observed. The time to 95 % agglutination was recorded. For these assays, a cutoff of 150 seconds was used because the 4 µL spots begin to dry.

#### Sperm surface galectin depletion

Sperm were washed in Multipurpose Handling Medium containing 100 mM lactose or asialofetuin (Sigma, Cat #: A4781), or 10 uM Olitigaltin (MedChem Express, Cat#: HY-19940) three times. Then, sperm were resuspended in fresh media. In rescue assays, after galectin depletion, either sterile filtered seminal plasma from the same donor (2.85 µL per million sperm), or recombinant Galectin-3 (10 µg per million sperm) and recombinant semenogelin-2 (100 pg per million sperm) were added to sperm cultures. Recombinant Galectin and semenogelin-2 were pre-incubated for 10 min at RT to pre-form complexes before addition to sperm cultures.

#### Measurement of sperm motility

Sperm motility was quantified using CASA software and specialized microscope slides with a shallow chamber depth. At least 300 sperm were measured per condition to determine a percent motility per time point.

#### Whole sperm ELISA

1.5×10^5 washed sperm in 50 μL of MHM were added to wells of a clear polystyrene Corning® ELISA 96-well microplate. The plate was incubated overnight, spun down at 300g, and the supernatant was discarded before the plate was air dried for 2 hrs at 37 °C. A standard curve of HCA doses were added in duplicate, and the remainder of the assay was performed using goat anti-human HRP secondary antibodies. The HRP-colorimetric signal was read at 450nm on a BioTek Synergy HTX multi-mode plate reader (Agilent, Santa Clara, CA, USA).

#### SDS PAGE and western blots

Sperm lysates were boiled in SDS containing 10% betamercaptoethanol (BME). Samples were loaded onto a 4–12% gradient polyacrylamide gel, and after electrophoresis, proteins were transferred to PVDF membranes. Loading control gels were stained in Coomassie dye to visualize protein bands. The blots were blocked with 5% powdered milk and washed with 0.1% TBS-T. All primary antibodies were used at a concentration of 10 µg/mL; anti-mouse (Santa Cruz Cat# sc-2005, RRID: AB_631736, Santa Cruz Biotechnology, Santa Cruz, CA, USA) and anti-rat (Abcam Cat# ab97057, RRID: AB_10680316, Cambridge, MA, USA) secondary antibodies were used at a 1:20,000 dilution, and the anti-human secondary antibody (Abcam Cat# ab6858, RRID: AB_955433) was used at a 1:5,000 dilution. Blots were developed in ECL substrate (ThermoFisher Scientific, Waltham, MA, USA). For fluorescent blots, Licor anti-human and anti-rat secondaries were used at dilutions described in manufacturer protocols.

#### On blot beta elimination

Blots were subjected to beta elimination by incubation of the PVDF membranes in 0.055 M NaOH for 16 hours at 37 °C prior to staining, then washed three times in TBS prior to staining.

#### In-gel digest of SDS PAGE

Samples corresponding to SDS-PAGE gel bands of <10 kDa, 10– 15 kDa, 15–20 kDa, and 20–25 kDa were excised and cut into small pieces. Gel pieces were subjected to reduction with 10 mM dithiothreitol (DTT) at 37 °C overnight, followed by alkylation with 50 mM iodoacetamide (IAA) at room temperature for 45 min in the dark. After washing with 50 mM ammonium bicarbonate and drying, the samples were digested with 50 ng of trypsin at 37 °C overnight. The resulting tryptic peptides were desalted using C18 ZipTips prior to LC– MS/MS analysis.

#### Immunopurification of HCA binders followed by mass spectrometry

HCA binders were isolated from sperm and seminal plasma by affinity chromatography using HCA-Protein G resin. These HCA-purified proteins were then analyzed by nano ultra-high-performance liquid chromatography-tandem mass spectrometry (nanoUPLC-MS and MS/MS) with nanoUPLC-MS/MS on an Orbitrap Fusion Lumos Tribrid mass spectrometer (Thermo Fisher Scientific, Waltham, MA, USA) coupled with an ACQUITY UPLC M-Class system (Waters Corp., Milford, MA, USA) via a TriVersa NanoMate (Advion, Ithaca, NY, USA). Detailed methods for the sample preparation and mass spectrometry data acquisition and analysis are provided in the supplementary material.

#### Mass spectrometry of seminal plasma O-glycopeptidome

Seminal plasma was prepared as previously described (43). Samples were analyzed by online nanoflow liquid chromatography-tandem mass spectrometry using an Orbitrap Eclipse Tribrid mass spectrometer (Thermo Fisher Scientific) coupled to a Dionex UltiMate 3000 HPLC instrument (Thermo Fisher Scientific). Data were acquired in data-dependent acquisition (DDA) mode, with MS/MS fragmentation performed via high-energy collisional dissociation (HCD) and additional electron transfer dissociation (ETD) fragmentation. Detailed methods for the sample preparation and mass spectrometry data acquisition and analysis are provided in the supplementary material.

#### HCA glycan array

A printed glycan microarray (CatchAll Glycan array, Cat #: 10616-8S, ZBioTech) was first blocked with blocking buffer (500 mg BSA in 50 mL PBST) for 1 hour and then incubated with HCA for 1 hour. Following incubation, the slides were washed four times with PBST, four times with PBS, and four times with deionized water, and then dried using a microarray slide spinner. The arrays were subsequently incubated with Alexa Fluor 647-conjugated anti-human IgG secondary antibody for 1 h at room temperature in a dark, humid chamber with gentle rocking. After secondary antibody incubation, the slides were washed sequentially with PBST, PBS, and deionized water prior to scanning. An image of bound fluorescence was then obtained using a microarray scanner (GenePix 4000 B, Molecular devices). Integrated spot intensities were acquired using Imagene software (GenePix Pro 7).

#### Statistical analysis

Continuous variables with >2 conditions were analyzed by analysis of variance (ANOVA) or mixed-model analysis. A significant effect was followed by Tukey or Holm-Šídák multiple-comparison test. Variables were tested for normality by the Shapiro-Wilk test. For repeated measures ANOVA, the equality of variances of the pairwise differences between within-subject conditions were assessed through tests of sphericity (Greenhouse-Geisser epsilon and Huynh-Feldt epsilon). Statistical significance was assumed when p<.05. Data analysis and graphing were performed with Prism, Version 9.4.1 610 (GraphPad Software, Inc, La Jolla, CA) software.

## Results

### Immunopurification and mass spectrometry indicate semenogelin is a component of the HCA antigen in both seminal plasma and sperm lysates

To identify the HCA binding partner, seminal plasma was used in initial studies as the antigen source because there were reports in the literature that seminal plasma was an abundant source of CD52, initially believed to be the HCA epitope. HCA was immobilized on a Protein G column, seminal plasma was added to the column, nonbinding elements were washed off, and the molecules that remained bound to HCA were eluted in low pH. The HCA binding partners were characterized by mass spectrometry. Abundant semenogelin-derived peptides were found in the purified material **(Table 1)**. Some minor contaminants were observed in the mass spectrometry analysis but at much lower levels than semenogelin **(Table 1)**. Notably, the CD52 peptide was not detected.

**Table 1:** Peptides assigned by mass spectrometry from HCA immuno-purification.

| **SOURCE: LIQUEFIED SEMINAL PLASMA** |  |  |  |  |  |
| --- | --- | --- | --- | --- | --- |
| **<u>PROTEIN OF ORIGIN</u>** | **<u>MW</u>**<br>**<u>(kDa)</u>** | **<u>GENE</u>** | **<u>INTENSITY</u>** | **<u>BAND</u>**<br>**<u>(kDa)</u>** | **<u>COVERAGE</u>**<br>**<u>(%)</u>** |
| **Semenogelin-2** | **71-76** | **SEMG2** | **1108815332.5** | **15-20** | **38.49** |
| **Semenogelin-1** | **52-55** | **SEMG1** | **984795391.9** | **15-20** | **48.48** |
| Trypsin | 23 | TRYP PIG | 101988673.3 | 15-20 | 7.79 |
| Keratin Type I cytoskeletal 9 | 62 | K1C9 | 43376886.9 | 15-20 | 28.41 |
| Keratin Type II cytoskeletal 1 | 66-70 | K2C1 | 39385849.4 | 15-20 | 34.32 |
| Keratin type I cytoskeletal 10 | 40-55 | K1C10 | 28068469.9 | 15-20 | 18.84 |
| Keratin type II cytoskeletal 2 epidermal | 65 | K22E | 17521793.0 | 15-20 | 20.66 |
| Antileukoproteinase | 14-18 | SLPI | 14100308.4 | 15-20 | 25.76 |
| Tyrosine-protein phosphatase non receptor type 13 | 277 | PTN13 | 10091359.2 | 15-20 | 0.68 |
| Keratin type II cytoskeletal 5 | 62 | K2C5 | 5887518.0 | 15-20 | 3.56 |
| Keratin type I cytoskeletal 14 | 52 | K1C14 | 2534521.4 | 15-20 | 9.75 |
| Human Immunoglobulin Heavy chain constant gamma 1 | 44 | IGHG1 | 1715379.7 | 15-20 | 13.64 |
| Keratinocyte proline rich protein | 64 | KPRP | 1500509.7 | 15-20 | 3.28 |
| **SOURCE: SPERM LYSATES** |  |  |  |  |  |
| **<u>PROTEIN OF ORIGIN</u>** | **<u>MW</u>** | **<u>GENE</u>** | **<u>INTENSITY</u>** | **<u>BAND</u>**<br>**<u>(kDa)</u>** | **<u>COVERAGE</u>**<br>**<u>(%)</u>** |
| Annexin A1;Annexin Ac2-26 | 40 | ANXA1 | 19848000 | 15-20 | 19.5 |
| Anosmin-1 | 76 | ANOS1 | 700820 | 10-15 | 1.6 |
| **Galectin-3** | **26** | **LGALS3** | **1719000** | **15-20** | **5.6** |
|  |  |  | 12004000 | 20-25 | 8.8 |
| Trypsin-3 | 25-27 | PRSS3 | 3505000 | 10-15 | 4.3 |
| **Semenogelin-2;Semenogelin-1;Alpha-inhibin-92;Alpha-inhibin-31;Seminal basic protein** | **52-55**<br>**71-76** | **SEMG1**<br>**SEMG2** | **1401800** | **15-20** | **3.8** |
| Lysozyme-like protein 2;Lysozyme-like protein 1 | 17; 12-21 | LYZ1,2 | 1043400 | 15-20 | 8.1 |
| Reticulon-4 receptor-like 1 | 49 | RTN4RL1 | 1.1322E + 10 | 15-20 | 2.9 |
| Ly6/PLAUR domain-containing protein 4 | 27 | LYPD4 | 3314300 | 15-20 | 4.1 |
| Sperm acrosome-associated protein 5 | 18 | SPACA5 | 10504000 | 15-20 | 4.4 |
| Protein S100-A8 | 11 | S100A7A | 12935000 | 15-20 | 11.4 |
| Desmocollin-1 | 100-110 | DSC1 | 1778800 | 10-15 | 1.5 |
| Solute carrier family 2, facilitated glucose transporter member 3;Solute carrier family 2, facilitated glucose transporter member 14 | 54 | SLC2A3 | 22710000 | 10-15 | 1.8 |
| Suprabasin | 60-65 | SBSN | 952160 | 15-20 | 6.1 |

This raised a question of whether HCA bound a different epitope in seminal plasma and on the surface of sperm. To answer this, sperm lysates were prepared by isolating motile sperm from several donors and pooling them to obtain sufficient amounts of protein. Sperm were lysed in 1% NP40 with sonication to gently disrupt cell membranes. Dot blots were performed to confirm the presence of CD52 (stained with Campath) and the HCA epitope prior to purification **(Supplementary Figure 1**). Immunoprecipitation was performed with a protein G column that contained either HCA, Campath, or a human isotype control antibody (VRC01, an anti-HIV antibody). The samples were run on SDS-PAGE and digested in-gel prior to mass spectrometry gel analysis. Prior reports in the literature have reported that CD52 does not stain with Coomassie dye, due its small size and limited number of amino acids that can bind the dye. Accordingly, small sections were cut from the gel for <10 KDa, 10-15 kDa, 15 - 20 kDa, and 20-25 Kda (containing the light chain of the antibody used for the IP), to capture CD52 that may not have reacted with the dye. Prior data from our lab suggest that even in the absence of a band by Coomassie stain, CD52 is present by western blot staining of the same gel, observed at a relative molecular weight of ∼20 kDa (**Figure 1**).

**Figure 1:**
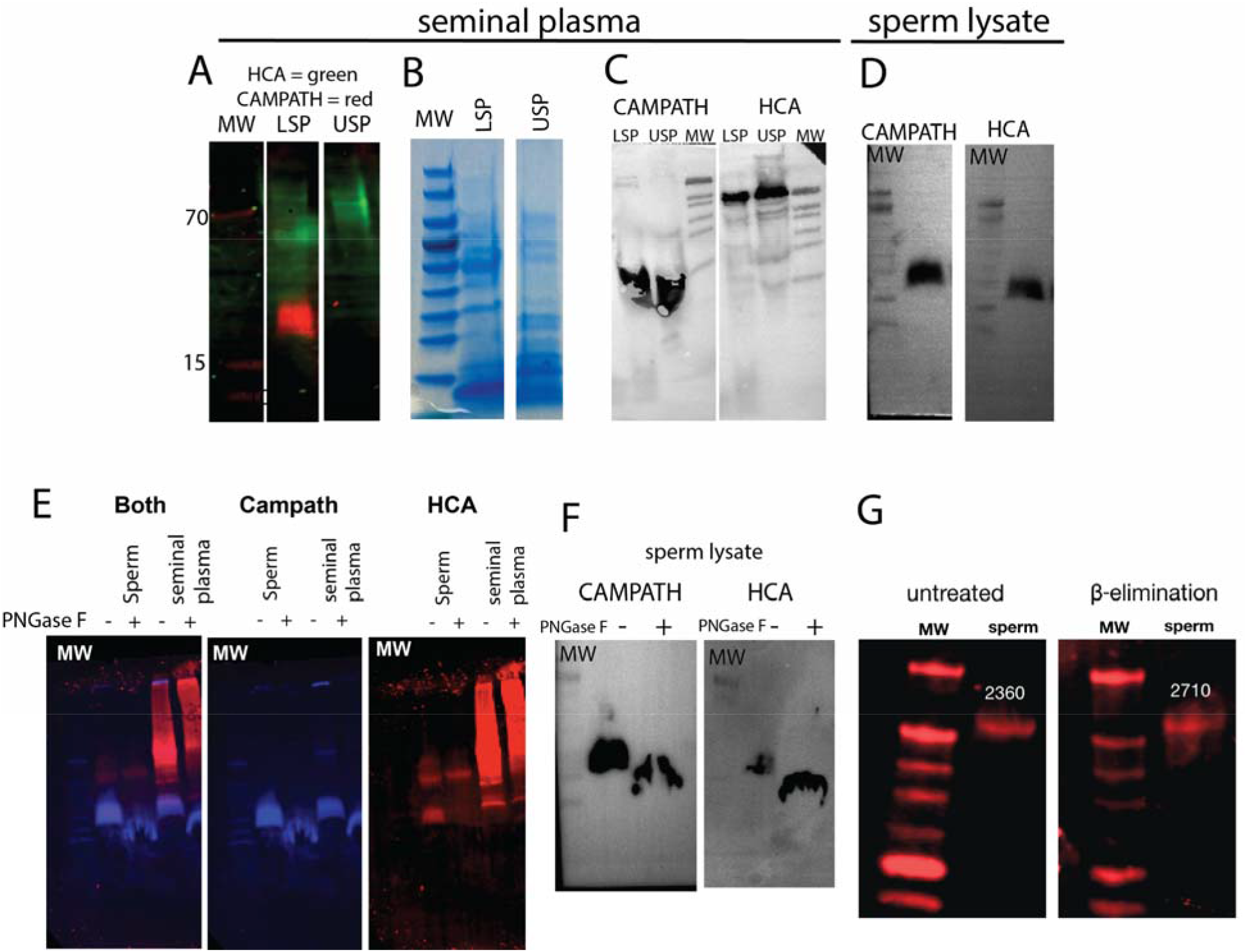
Western blots demonstrate that HCA binds a larger protein relative to CAMPATH (anti-CD52) in seminal plasma, and that N-linked deglycosylation impacts the molecular weight of the glycoprotein recognized by HCA. A) Fluorescent Western blot of liquified (LSP) and unliquefied seminal plasma (USP). Red color represents CAMPATH (anti-CD52 antibody) binding and green represents HCA binding. B) SDS-PAGE containing both USP and LSP as a loading control for the western blots. C) Chemiluminescent western blot of unliquefied and liquified seminal plasma from a Tris-tricine gel to further separate smaller peptides. D) Chemiluminescent western blot of sperm lysates. E) Fluorescent Western blot with sperm lysates and seminal plasma. Plus indicates presence of PNGase F and minus is a negative control. Blue represents CAMPATH (anti-CD52 antibody) staining and red represents HCA staining. F) Chemiluminescent blot of sperm lysates. Plus indicate PNGase F and minus indicates negative control. G) An untreated and beta elimination western blot of sperm lysate, with HCA staining in red. Number indicates quantification of observed band.

These sperm immunoprecipitation experiments produced lists of several hundred proteins. These lists of proteins included both nonspecific interactors and parts of the HCA binding complex. Notably, CD52 was not identified from the CAMPATH-purified material or the HCA purified material. This may be due to CD52’s high degree of glycosylation and small size, which may hinder its release from an SDS-PAGE gel through in-gel trypsin digest sample preparation; in addition, we cannot account for potential uncharacterized, extensive glycan modifications (a large *N*-linked glycan, some *O*-linked glycosites, and a GPI anchor) that may preclude identification of the CD52 peptide in a database search.

To narrow down the proteins identified from the HCA purified material to a higher confidence, shorter list of potential HCA binders, the immunoprecipitation data were filtered to remove potential contaminants identified by the Byonic software, proteins for which the statistical significance value q >0.05, and proteins for which intensity was zero. Next, we used internal controls to further filter proteins as nonspecific. First, the proteins identified in both HCA-purified material and VRC01(isotype control)-purified material were classified as nonspecific binders and removed. Next, common contaminants in affinity purifications, as defined by appearing in over 100 experiments in the CRAPome database, were removed (*12*). Finally, since it is known that HCA’s epitope is a sperm surface protein that is also secreted into seminal plasma, the list of proteins of interest was enriched for secreted proteins using the subcellularviz tool (*13*). The resulting list of the proteins of interest from sperm lysate is summarized in Table 1. Of these proteins, the ones identified from the 15-20 Kda band that was cut from the gel are of particular interest since HCA staining is observed on western blots of sperm lysate in that apparent molecular weight range. This workflow is summarized in **Supplementary Figure 2.** These proteins include ANXA1, LGALS3, SEMG1, SEMG2, LYZ1, LYZ2, RTN4RL1, LYPD4, SPACA5, S100A7A, SBSN. Notably, semenogelin-1 and −2 were identified in both the seminal plasma and the sperm lysate **(Table 1),** and there was a single semenogelin-derived tryptic peptide (ISYQSSSTEER) that was isolated from both the sperm lysates and seminal plasma, though this peptide alone did not reduce HCA activity by competition **(Supplementary Figure 3, Supplementary Figure 4).**

### Western blotting and deglycosylation studies implicate larger proteins as potential HCA binders

To replicate experiments from the literature, western blots of seminal plasma and sperm lysates were prepared, containing both unliquefied and liquefied seminal plasma to account for physiological liquefaction altering the seminal glycoproteome. We found that HCA binding localizes differently on western blots depending on the source of the antigen. In seminal plasma HCA recognizes a large size range of proteins, whereas in sperm lysates it recognizes proteins corresponding to a smaller apparent molecular weight **(Figure 1A, 1D, loading control is Figure 1B).** Campath, a CD52 specific monoclonal antibody, binds to similarly sized regions of the blot in both sperm lysates and liquefied seminal plasma **(Figure 1A, 1D).** We found that Campath does not bind to unliquefied seminal plasma, probably because un-liquefied seminal plasma does not contain soluble CD52, as previously described ((11), **Figure 1A**). Notably, HCA did bind un-liquefied seminal plasma on blot, supporting the mass spectrometry data indicating semenogelin as a HCA binding partner **(Figure 1A)**.

To investigate the role of *N*-linked glycans in the HCA antigen, we treated the liquefied seminal plasma and sperm lysates with PNGase F, a bacterial enzyme that cleaves *N*-linked glycans from proteins. For Campath staining, deglycosylation resulted in a gel shift of the CD52 band, which is to be expected and serves as a positive control for successful deglycosylation of the lysate **(Figure 1E).** Since Campath’s epitope is a peptide sequence and not reliant on the presence of the N-glycan, the staining intensity for CD52 remains similar in both sperm and seminal plasma blots **(Figure 1E).** For HCA staining, the impact of PNGase F treatment was variable depending on the antigen source. In the sperm lysates, HCA staining resulted in two bands, one of which overlapped with the Campath positive region, and one band that was entirely separate **(Figure 1F).** PNGase F treatment eliminated the staining for the Campath overlapping band, which may indicate that for the protein in that region that is HCA binding, the *N*-linked glycan is required for binding. The other sperm lysate band was not impacted by PNGase F treatment indicating that some proteins in sperm may bind to HCA independent of an N-glycan. For seminal plasma, HCA binding was seen across large regions of the blot as a smear, and it was unclear whether there was any gel shift or change in binding due to the removal of the N-glycans. Finally, to determine if HCA interacted with *O*-linked glycosites, sperm lysates were blotted and then subjected to on blot beta elimination prior to staining. The intensity of staining was unchanged after beta elimination, indicating that the binding site of HCA does not contain or is not impacted by the presence of an *O*-linked glycan **(Figure 1G).** Loading controls for sperm lysates are in **Supplementary Figure 5.**

Western blotting of these materials has given conflicting results, potentially due to several factors including heterogeneity in human semen samples, physiological liquefaction, complex glycosylation, or variation in secondary antibody sensitivities. Switching from chemoenzymatic blots to fluorescent blotting revealed additional binding sites of the antibody on the western blots. Accordingly, to study this particular material, other methods may be better suited, and these results are interpreted with caution and with several additional confirmatory experiments.

### In-gel digest Mass spectrometry of HCA-reactive regions of polyacrylamide gel reveals that semenogelin is present in regions of the gel that react with both HCA and CAMPATH on blots

To understand the potential confusion between CD52 and semenogelin derived peptides, which has been previously reported, we cut regions of an SDS gel of seminal plasma that reacted with Campath and HCA on western blots, digested the proteins in-gel, and assigned peptides by LC/MS-MS analysis **(Figure 2A)**. Next, we compared the assigned peptides from those regions to the peptides assigned to the HCA immuno-purified material. We found an overlap of three proteins in both HCA-reactive and Campath-reactive regions of the gel, and in the HCA-immuno purified material: semenogelin-1, semenogelin-2, and human IgG heavy chain (**Figure 2B**). Notably, CD52 was not identified in any sample indicating that it may not be detectable by in-gel digest LC/MS-MS. Semenogelin derived peptides may be responsible for some of the HCA staining that was attributed to CD52 in western blots reported in the literature. In addition, semenogelin derived peptides are present in several regions on gels and western blots, likely due to physiological cleavage by endogenous proteases during semen liquefaction. Semenogelin is a major component of seminal plasma, and its presence in high milligram quantities in seminal plasma may crowd out the presence of other lower abundance proteins in mass spectrometry of unpurified material. These data on seminal plasma indicate that conventional methods to determine antibody specificity, including western blotting, may fall short on such a highly glycosylated, endogenously cleaved, and abundant protein such as semenogelin.

**Figure 2:**
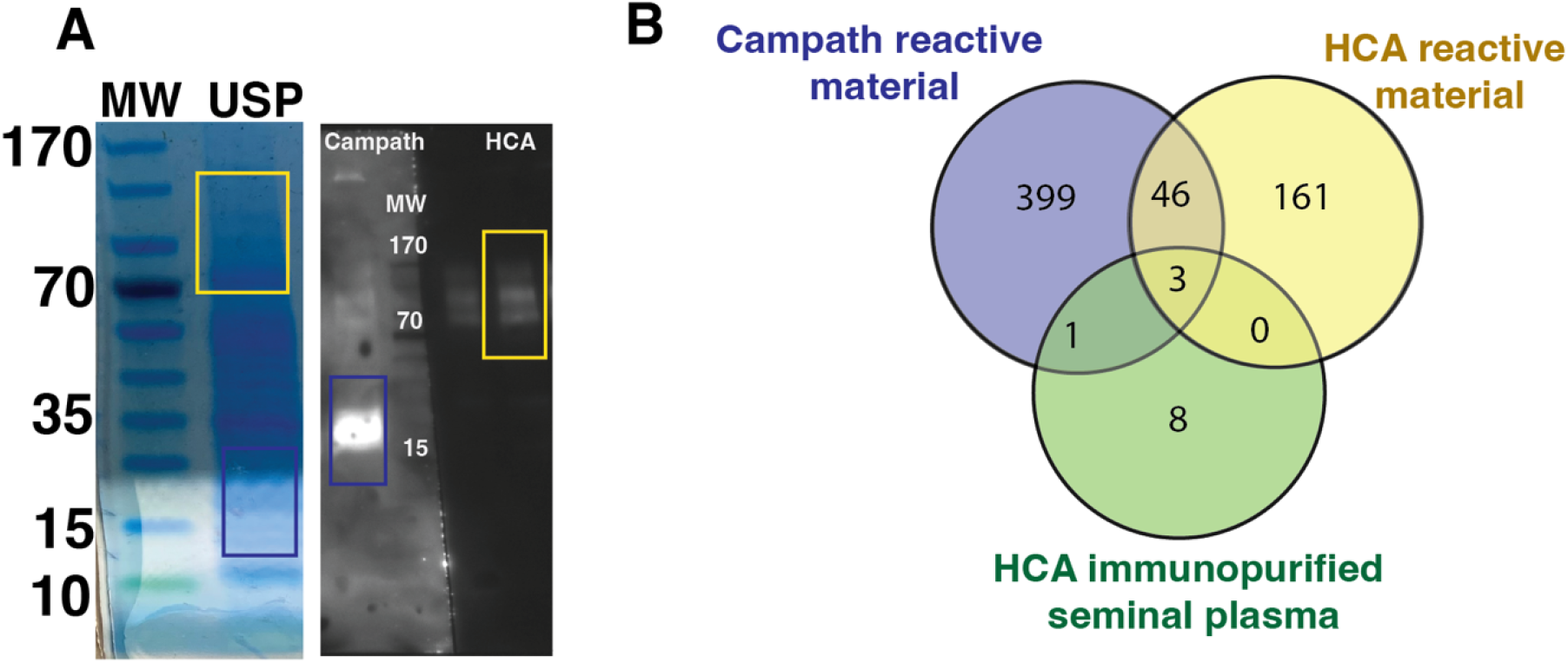
Comparison between HCA and CAMPATH reactive regions of gel and HCA immuno-purified seminal plasma. (A) SDS-PAGE and western blot analysis of unliquefied seminal plasma, indicating the gel regions excised for mass spectrometry analysis. The yellow box denotes the region corresponding to the HCA-reactive section of the Venn diagram, while the blue box denotes the region corresponding to the CAMPATH-reactive section. (B) Venn diagram illustrating the overlap in proteins identified by LC-MS/MS in HCA-purified material and in the HCA- and CAMPATH-reactive gel regions.

### HCA binding requires an *N*-linked glycan on the sperm surface, but does not demonstrate reactivity with any particular glycan on array

To determine if sperm surface N-glycans are part of the HCA epitope, intact sperm were exposed to PNGase F for 2 hours, fixed, then stained with HCA and analyzed by flow cytometry. We found that HCA staining was significantly reduced after removal of *N*-linked glycans on the surface of sperm **(Figure 3A).** This indicates that either the sperm epitope of HCA contains an *N*-linked glycan, or that the epitope remains attached to the surface of sperm via an *N*-linked glycan, or both. We also observed that enzymatic removal of *N*-linked glycans from sperm surface abrogates HCA-mediated agglutination (**Figure 3B).**

**Figure 3:**
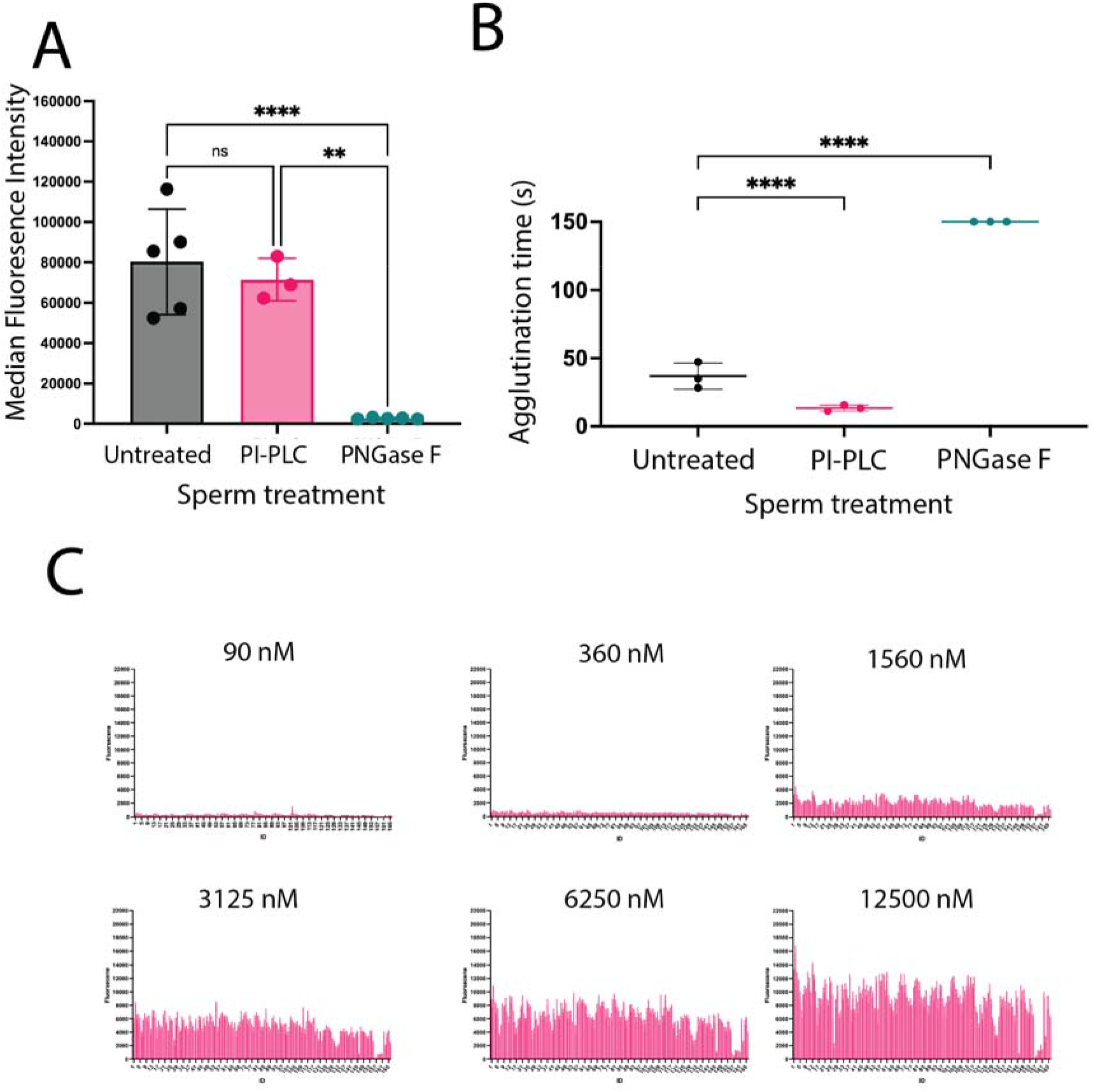
Flow cytometry of sperm demonstrates that HCA binding requires the presence of an N-linked glycan, but glycan array data demonstrate that HCA does not directly bind a specific glycan. A) Flow cytometry of sperm after HCA staining. Data represent the median fluorescent intensity of HCA binding to sperm. Sperm were treated with PNGase F or PI-PLC for 1 hour, then fixed, and finally stained with 5 ug per mL HCA, which corresponds to a 33 nanomolar concentration. B) Kinetic agglutination after treatment with PNGase F and PI-PLC. Points are the mean of three technical replicates. C) HCA binding to the CatchAll glycan array at varying concentrations. HCA was incubated on the array at the indicated concentrations, followed by staining with an anti-human IgG secondary antibody. Bars represent mean fluorescence intensity averaged across 4 technical replicates for each glycan structure on the array. Structures represented on the array are numbered on the X axis and described in Supplementary Figure 3. Statistics represent mixed model or 2-way ANOVA comparisons with post-hoc Holms-Sidak tests in Fig A and B.

To test whether this proposed HCA binding complex included a GPI anchored protein such as CD52, sperm were treated with PI-PLC prior to kinetic agglutination. Interestingly, removal of PI-PLC sensitive GPI anchored proteins enhanced HCA mediated agglutination but did not enhance binding **(Figure 3B),** suggesting several possibilities: 1) that the HCA epitope does not adhere to the sperm surface through a PI-PLC sensitive GPI anchor, 2) that the HCA epitope adheres via a GPI anchor that is resistant to PI-PLC mediated cleavage, or 3) removal of GPI anchors modifies sperm biology in a way that promotes sperm agglutination. Given that PI-PLC does not remove all GPI anchored proteins, these data cannot conclusively determine whether the HCA epitope complex is bound to sperm via a GPI anchor or not.

To determine which glycans HCA may directly interact with, we quantified the binding of HCA to a glycan array representing of several glycan classes **(Supplementary Figure 6)**. In this array, we observed broad nonspecific reactivity with several classes of glycans. We observed significant binding from 3.125 mM to 12.5 mM range (**Figure 3C**). However, the activity of HCA on the surface of sperm is observed in the nanomolar concentration range where binding was not observed. In addition, competition with the “top hit” (GlcNAc) on the glycan array did not impact HCA binding in a whole sperm ELISA leading us to believe the signal from these higher concentrations of antibody is largely nonspecific **(Supplementary Figure 7).** Accordingly, we hypothesize that the overall presentation of glycans in the array format may not recapitulate the native glycan configuration that exists on the cell surface, resulting in lower observed binding affinities on-array compared to the surface of sperm. In addition, we further hypothesize that avid presentation of glycoforms, such as in a dense lipid raft, or through repeat glycan moieties enhance the binding affinity of HCA to sperm glycans. These data raise the possibility that HCA does not directly bind a glycan but relies on the presence of an *N*-linked glycan for anchoring or adhesion of its epitope on the sperm surface.

### Competition with recombinant semenogelin proteins actually enhances HCA activity

We hypothesized that if HCA binds semenogelin, introducing recombinant semenogelin to the kinetic agglutination assay should reduce the binding of HCA to sperm and thus decrease sperm agglutination. Addition of the semenogelin alone to sperm did not induce agglutination in the absence of HCA. However, surprisingly, when we preincubated HCA with the semenogelins, then exposed these HCA immune complexes to sperm, agglutination was enhanced (eg. time to agglutination was decreased) **(Figure 4A).** This decrease was greater for semenogelin-2 than for semenogelin-1 **(Figure 4A).** This indicates that the sperm may directly bind semenogelin, providing a higher antigen density for HCA to bind to, resulting in faster agglutination **(Figure 4B)**.

**Figure 4:**
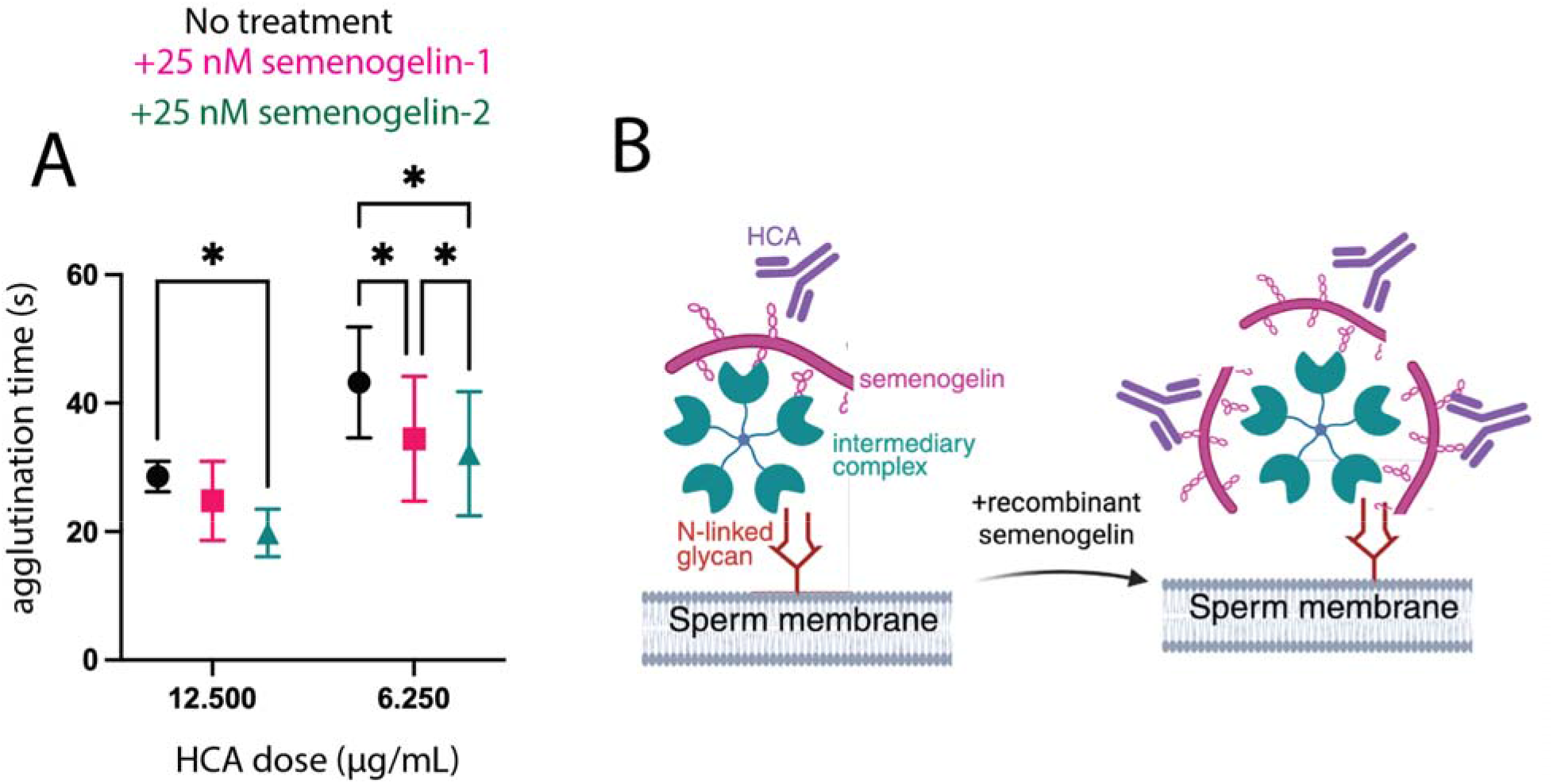
Recombinant semenogelin enhances HCA-mediated agglutination. A) Time to agglutination in the presence and absence of recombinant semenogelin-1 and semenogelin-2. B) schematic representing hypothesis that agglutination is enhanced because semenogelin adheres to the sperm surface and increases the density of the HCA target. Statistics represent mixed model or 2-way ANOVA comparisons with post-hoc Holms-Sidak tests.

**Figure 5:**
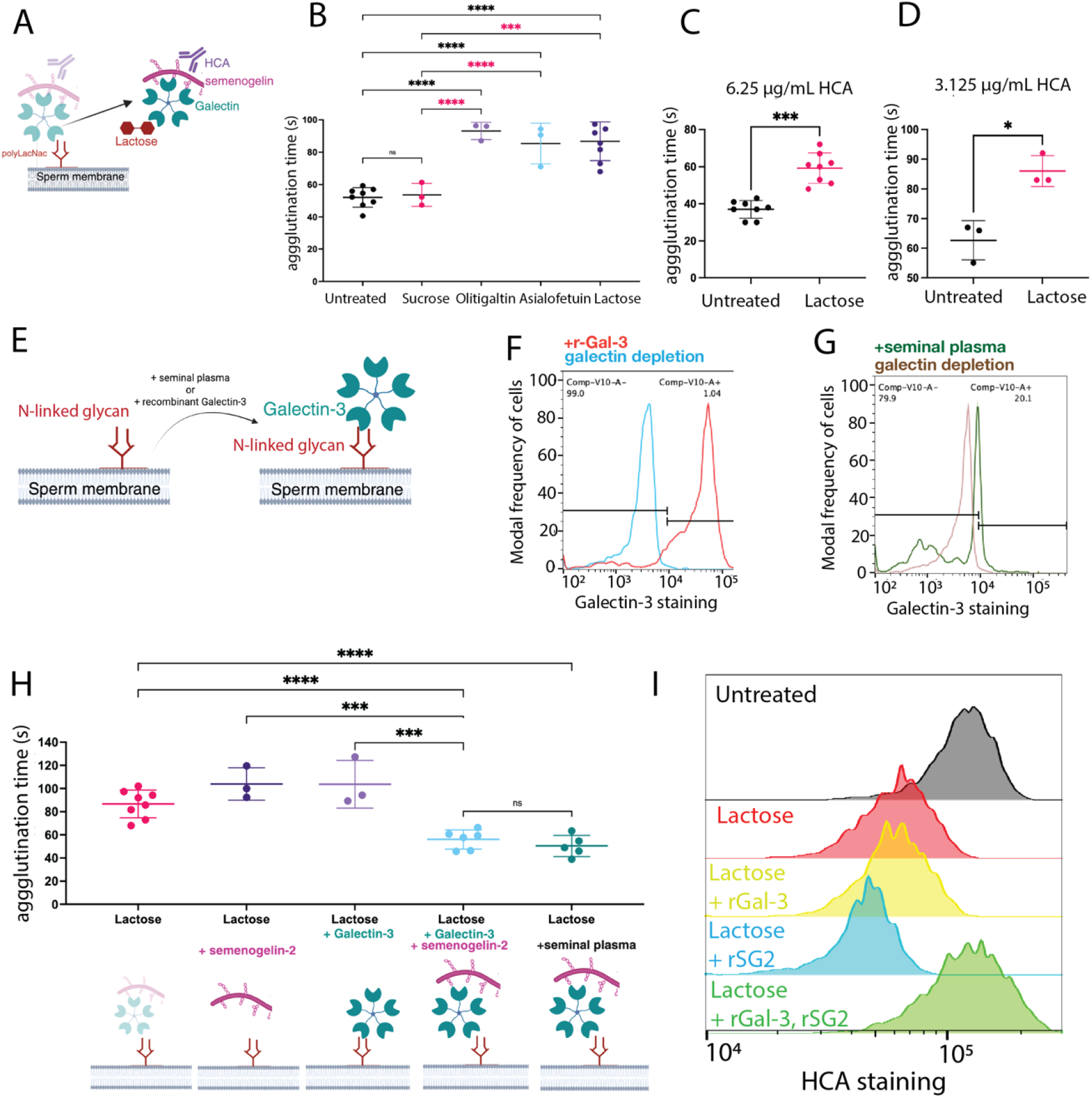
Depletion of galectin on the sperm surface reduces HCA-mediated agglutination activity; this can be rescued by addition of seminal plasma or recombinant galectin-3 and semenogelin complexes. A) Schematic describing galectin depletion on sperm surface by competitive inhibition. B) Time to agglutination in the presence of HCA (6.25 ug/ml) after sperm were treated with several different galectin inhibitors. No treatment and sucrose treatment were negative controls for galectin depletion. C) Time to agglutination in the presence of HCA (6.25ug/ml) after galectin depletion. D) Same as E, but with 3.125 ug/mL HCA. E) Schematic representing exogenous addition of Galectin-3 onto sperm surface after galectin depletion. F) Flow cytometry of sperm after galectin depletion followed by exogenous addition of recombinant Galectin-3. G) flow cytometry of sperm after galectin depletion and then exogenous addition of 10% seminal plasma. H) Time to agglutination in the presence of 6.25 ug/mL HCA after galectin depletion, and exogenous addition of recombinant galectin, semenogelin and/or seminal plasma. I) Flow cytometry of sperm after galectin depletion and addition of recombinant galectin-3 and semenogelin. Statistics represent mixed model or 2-way ANOVA comparisons with post-hoc Holms-Sidak tests.

### Semenogelins carry terminal LacNAc O-glycans

HCA-mediated agglutination is enhanced by exogenous addition of recombinant semenogelin proteins and is dependent on the presence of N-glycosylation on the sperm cell surface, despite HCA not binding strongly to any particular glycan structure. A framework that may explain this is as follows: HCA binds to semenogelin, which in turn binds to sperm via a sperm-surface N-glycan-dependent mechanism. An interaction between the heavily O-glycosylated SEMG proteins and a sperm-surface N-glycan would likely be mediated by a galectin, a class of proteins with carbohydrate-binding domains that have been shown to cross-link or anchor cells (17, 20, 26). As such, Galectin-3 (Gal-3), which was detected by mass spectrometry in the HCA pulldown, is a strong candidate for this role. Gal-3 is a glycan-binding protein that preferentially binds to terminal LacNAc glycan moieties, which are found on both N- and O-glycans (42). It exists as a pentamer when secreted, and has been shown to form heterogeneous cross-linking complexes with multivalent carbohydrates (42). It has been shown previously to be expressed in seminal fluid and sperm cells, as well as being found in our mass spectrometric analysis of sperm lysate immunoprecipitation **(29, 31, Table 1).**

In order for Gal-3 to anchor the SEMG/HCA complex onto the sperm cell surface, terminal LacNAc moieties must be present on one or both semenogelins. A prior glycoproteomic study of seminal plasma revealed 17 O-glycosites on SEMG1 and 20 glycosites on SEMG2 (34). As such, we considered that *O-*linked glycans on the SEMG proteins may act as binding partners for gal-3 during agglutination. However, the previous investigation searched only for 6 of the hundreds of possible human O-glycan structures, none of which contained a LacNAc moiety, the preferential binding partner of gal-3. Additionally, the aforementioned preparation made use of Zic-HILIC enrichment, which is not ideal for O-glycoproteomic analyses due to its bias towards glycopeptides with high glycan-to-peptide ratios by mass (i.e. N-glycans) (34). Despite these shortcomings, semenogelin-derived peptides in seminal plasma still seemed strong candidates for harboring *O-*linked LacNAc moieties. To investigate this, we conducted an O-glycopeptidomic analysis of endogenously generated SEMG peptides using a modified GlycoFASP protocol (43). In short, liquified human seminal plasma was placed on a 30kDa or 10 kDa MWCO filter, and peptides generated from endogenous proteases such as PSA were collected via centrifugation. To generate small peptides amenable to LC-MS/MS analysis, we digested the 30 kDa sample with the O-glycoprotease IMPa, which cleaves peptides N-terminal to an O-glycosite, and with trypsin. Peptides were first identified in the MS using HCD fragmentation to identify sequences. Parent ions that generated spectra containing oxonium ions from glycan fragmentation were then reselected for EtD or EThcD fragmentation; a softer dissociation technique that retains the glycan on the peptide backbone.

In our analysis, we identified many O-glycosites on both SEMG proteins, including those consistent with terminal LacNAc moieties **(Supplementary Figure 8).** However, standard proteomics search tools often struggle to correctly identify which O-glycosite is occupied in a peptide with multiple potential O-glycosites. To account for this, manual validation of glycopeptide spectra is required to confirm which glycosites harbor terminal LacNAc moieties. To this end, we manually validated all O-glycopeptides identified by the Byonic search software with scores above 150. Sequences were considered validated if the intact naked peptide ion and at least 3 b- and y-ions were observed in HCD spectra. Glycosites were validated from either EtD or EThcD spectra that contained at least 1 c- and z-ion on either side of an occupied glycosite. With this we identified 17 O-glycosites on SEMG1 and 23 O-glycosites on SEMG2. Of the O-glycosites identified in this analysis, 9 of the O-glycosites from SEMG1 and 7 of the O-glycosites from SEMG2 were identified in the previous analysis. Some of the O-glycosites found in the previous study were not found here, likely due to differences in sample preparation. Our analysis aimed to collect endogenously generated peptides and did not use any glycopeptide- or glycan-specific enrichment, whereas the previous exploration digested the entire seminal fluid proteome before performing glycan-specific enrichment. Of the O-glycosites identified in our analysis, 6 from SEMG1 and 2 from SEMG2 harbored glycans consistent with terminal LacNAc moieties. Additionally, 5 glycopeptides from SEMG1 and 10 glycopeptides from SEMG2 were identified in an O-glycoproteomic analysis of HCA immunoprecipitated seminal fluid and sperm lysate. Interestingly, 2 O-glycosites from SEMG1 and 1 O-glycosite from SEMG2 that we identified in the HCA pulldown were found to harbor terminal LacNAc moieties in our targeted O-glycoproteomic approach **(Tables 2 and 3).**

**Table 2:**
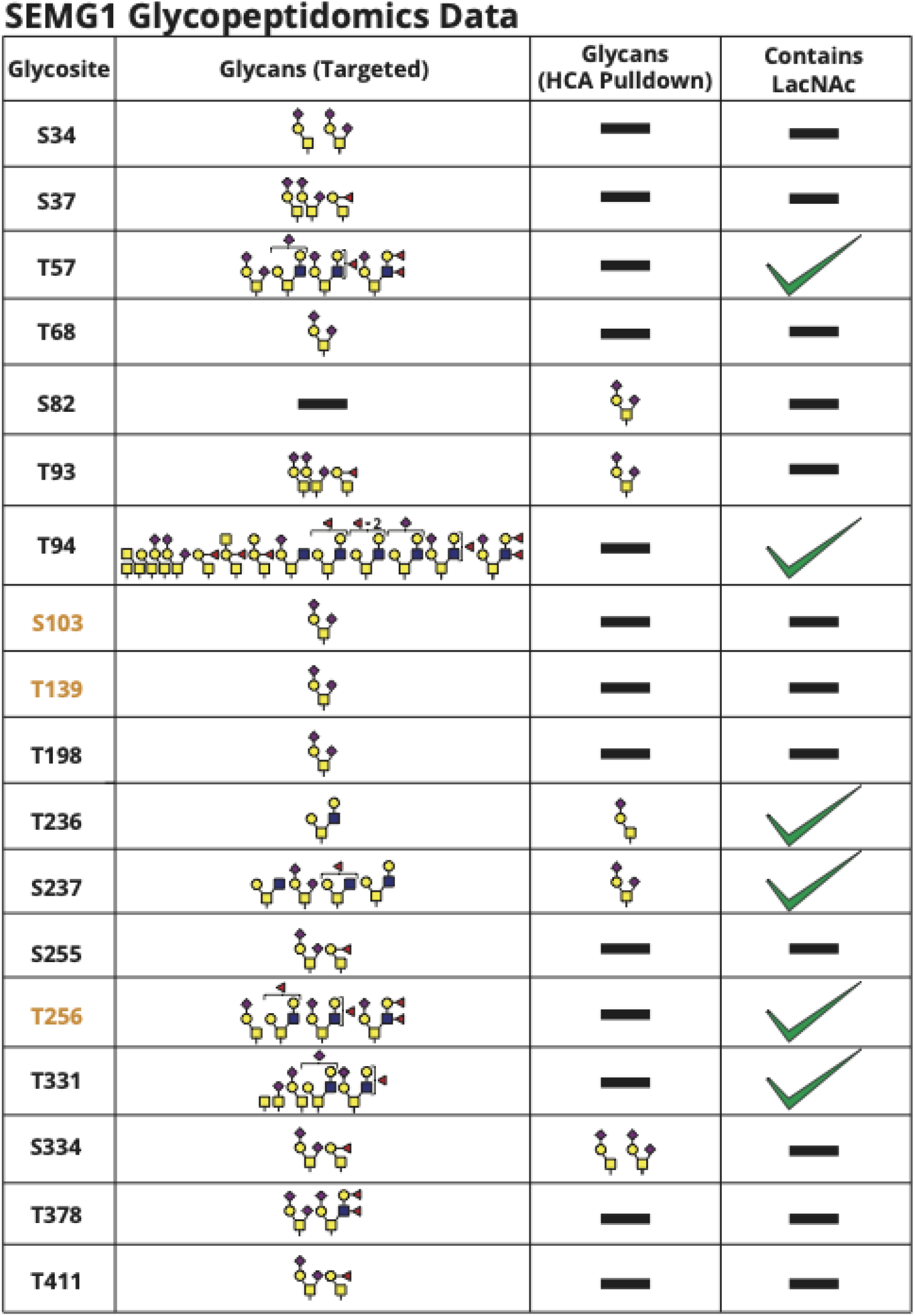
SEMG1 Glycopeptidomics Data and HCA pulldown Glycoproteomics data.

**Table 3:**
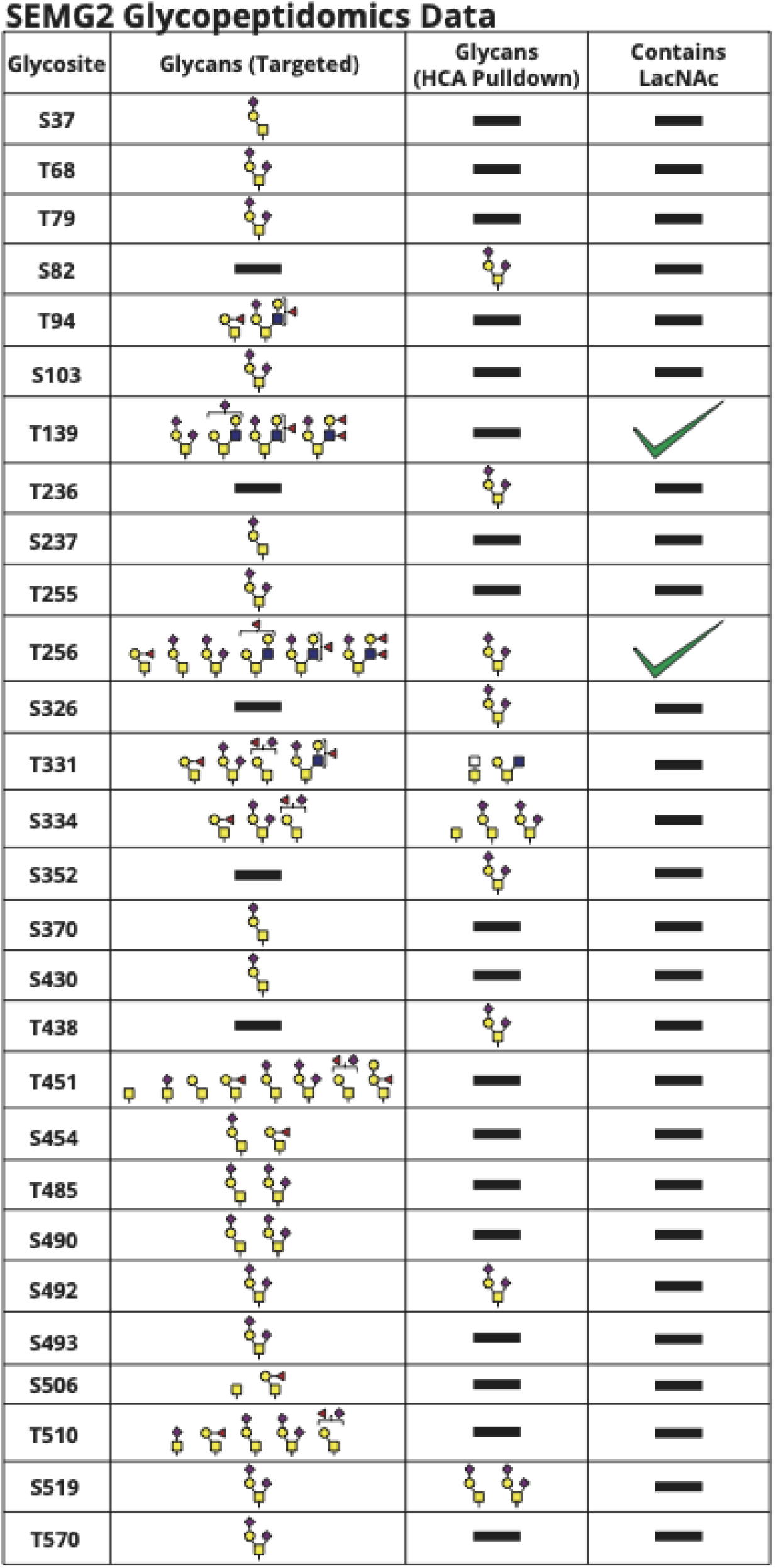
SEMG2 glycopeptidomics data and HCA pulldown glycoproteomics data.

It is important to note that HCA and Gal-3 may not need to bind to the same region of SEMG proteins to form a protein complex. There is, however, an abundance of O-glycans, including LacNAc moieties, found on the semenogelins from our targeted O-glycoproteomics, along with a plethora of SEMG peptides isolated from HCA purification. These results, when taken together, suggest that SEMG proteins can bind both HCA and Gal-3, possibly through multivalent binding. Overall, this lends support to the idea that SEMG O-glycopeptides are responsible for bridging Gal-3 and HCA, allowing HCA-mediated agglutination.

### HCA-mediated agglutination is reduced by galectin depletion on the surface of sperm and can be rescued by adding seminal plasma or recombinant galectin-3 and semenogelin complexes to sperm cultures

To test the model that Gal-3 could be a connecting link between sperm *N*-linked glycans and the docking of the HCA epitope (semenogelin) on the sperm surface, we sought to remove the galectin/semenogelin complex from the surface of sperm and assess HCA agglutination function. Prior reports in the literature indicate that soluble LacNac and lactose treatments can deplete sperm of surface Galectin-3 (*17*). Accordingly, we pursued several agglutination experiments after depleting sperm surface galectins with soluble lactose, asialofetuin and olitigaltin (a commercially available cancer drug competitive inhibitor of galectin) to determine how galectin depletion may impact HCA-mediated agglutination **(Figure 4A).** To confirm that galectin depletion did not impact agglutination by reducing sperm motility, we measured sperm motility and found that galectin-depletion sperm treatments used in this set of experiments did not impact sperm motility **(Supplementary Figure 9A, 9B, 9C)**. We found that washing sperm with competitive galectin inhibitors significantly increased the time to HCA-mediated agglutination **(Figure 4B)**. Time to agglutination was increased by 1.6-fold by lactose and asialofetuin treatment, and by 1.7-fold by olitigaltin treatment **(Figure 4B)**. The lactose mediated increase in agglutination time was retained across two different HCA concentrations, at both 3.125 and 6.25 µg/mL **(Figure 4C**, **4D).**

A key issue throughout this study of the HCA antigen has been the infeasibility of applying genetic tools to ex-vivo human sperm cultures. Typically, to study antibody binding and conclusively determine an antigen, cells that both express the protein of interest and can be grown in culture are used to generate matched negative control knockout cells where protein expression is absent. Given the lack of methodology to modify protein expression in ex vivo human sperm genetically, conventional molecular biology rescue experiments for HCA function were out of reach. However, because galectin depletion does not require genetic modification of the sperm, is compatible with the limited time frame of ex vivo sperm cultures (<5 hours) and demonstrates a reduction in HCA-mediated agglutination function, it can serve as an alternative way to perform molecular rescue experiments. To do so, we depleted sperm surface galectins with lactose as before and exogenously added different predicted components of the HCA binding complex **(Figure 4E).** We first added back sterile filtered seminal plasma as a positive control for the rescue (at a 1:10 ratio to the sperm culture), which is a complex mixture containing both soluble galectins and semenogelin-derived peptides, and is known to contain the soluble HCA epitope as determined by western blot **(Figure 1).** We found that adding seminal plasma into the cultures restored sperm-surface Gal-3 by flow cytometry (**Figure 4G).** As expected, adding the seminal plasma restored HCA-mediated agglutination speed back to a comparable level to the untreated sperm (**Figure 4H**). Next, we added back recombinant human Galectin-3 (rGal-3) and recombinant human semenogelin-2 (rSG2) to the cultures separately. We confirmed binding of the rGal-3 onto the sperm by flow cytometry **(Figure 4F).** rGal-3 and rSG2 alone did not rescue HCA-mediated agglutination (**Figure 4H)**. Finally, we precomplexed rGal-3 and rSG2 for 5 minutes, then added the complexes to sperm cultures prior to agglutination. This rescued HCA-mediated agglutination back to a comparable speed to the untreated control **(Figure 4H).** Finally, we stained sperm with HCA after galectin depletion and rescue, finding that HCA staining by flow cytometry followed the same pattern we had seen with HCA-mediated agglutination **(Figure 4I)**.

## Discussion

These data suggest that CD52 may not be required for HCA binding. In this work, we first purified the HCA epitope from liquefied seminal plasma and sperm and characterized this HCA-purified material by mass spectrometry. From both seminal plasma and sperm, semenogelin-derived peptides were isolated and CD52 was not identified. Notably, CD52 was not identified in samples prepared by immunoprecipitation using CAMPATH (anti-CD52 antibody) and an in-gel digest prior to LC/MS-MS analysis, pointing to the need for highly specialized purification and detection methods to detect GPI anchored CD52 due to its small size, heterogenous GPI anchor, and high degree of glycosylation. There was some overlap in the proteins identified by mass spectrometry for CAMPATH and HCA immunopurified samples, and semenogelins appeared in both. It is possible that the *N*-linked glycan of CD52 anchors a semenogelin-containing complex to the surface of sperm.

Next, we found that HCA interacts with seminal plasma proteins larger than CD52 that present as a large smear on western blots, which is different from previously reported data (*1*, *14*). Semenogelin is highly glycosylated and endogenously cleaved as part of its physiological function of semen liquefaction and thus can appear as a smear at various apparent molecular weights on gels and western blots (*11*, *18*). The fact that HCA stained a large spread of protein sizes in western blots of seminal plasma fits the profile of semenogelin which varies in size due to physiological cleavage by endogenous proteases. In addition, HCA binding was observed in unliquefied seminal plasma samples that reportedly do not contain CD52 (11) and these samples did not bind Campath (anti-CD52 antibody) in our experiments. Prior literature has reported that HCA cross-reacted with material that bound S19 (a different antisperm antibody) on western blots, which was identified as CD52 by Edman degradation (*9*). Edman degradation, which is not suited for complex mixtures of proteins such as those derived by immuno-precipitation of biological materials, may not have identified all peptides in the material. This is particularly important since it has been observed that semenogelin and CD52 run at identical apparent molecular weights on gels and can co-purify in immunoprecipitation (11). Additionally, prior literature reported that HCA directly bound a polyLacNAc motif by thin layer chromatography, but the study lacked any characterization of the placenta derived glycolipid standards used to assign the HCA binding site as polyLacNAc (5). To further investigate the possibility that HCA binds semenogelin rather than CD52, we determined which proteins were present in the regions of a gel that reacted with both CD52 and HCA on blots, finding that semenogelin-derived peptides were present in both. The unfortunate fact that both CD52 and semenogelin-derived peptides on the surface of sperm both copurify and run at identical relative molecular mass as glycosylated smears on SDS PAGE (11) and that biochemical methods in prior literature were limited (Edman degradation and western blotting) and overinterpreted can explain why the field coalesced on CD52 as the HCA antigen without definitive evidence (1, 5, 9, 11). Our findings here consistently suggest that semenogelin may be the HCA target, and that CD52 is not required for HCA binding alone.

Prior literature has reported consistent HCA binding on western blots at a smaller apparent molecular weight of approximately 15 - 20 Kda (1). This discrepancy can be attributed to choice of staining technique and key differences in lysate preparation. Prior literature did not use fluorescent staining which has a much higher sensitivity than chemoenzymatic staining and did not test unliquefied seminal plasma. A key difference between the sperm lysate blots described in this study and western blots of sperm lysate in the literature are in the minute details of sample preparation. In Baldeon et. al. deglycosylated sperm lysates were prepared by treating intact sperm with PNGase F prior to lysis (1). In this study, deglycosylated sperm lysates were prepared by lysing sperm first then deglycosylating with PNGase F. Given that semenogelin may adhere to sperm through a *N-*linked glycan-dependent interaction on the sperm surface, whether deglycosylation is performed before or after lysis can change the absence of presence of HCA staining from the same starting material.

To investigate the glycan binding profile of HCA, we treated intact sperm with PNGase F to remove all *N*-linked glycans and found that removal of *N-*linked glycans abrogated HCA binding. However, when probing HCA binding to glycans directly in an array format, we found no evidence of a specific interaction between HCA and a particular glycan on the array. In addition, deglycosylation by PNGase F (to remove *N*-linked glycans) or on-blot beta elimination (to remove *O*-linked glycans) did not consistently abrogate HCA binding on western blot. These conflicting data led us to the idea that HCA binding on the surface of sperm requires an *N*-linked glycan, but that HCA did not directly bind an *N*-linked glycan.

One potential model that would fit these data is that an *N-*linked glycan anchors an intermediary multivalent glycan-binding protein that also binds to semenogelin. It is possible that complex N-glycans on sperm interact with a multivalent glycan binding protein, such those in the galectin family, which would allow the multimeric galectin to serve as an anchoring site for semenogelin-derived glycopeptides that could bind HCA directly **(Supplementary Figure 10, Supplementary Table 1).** Analysis of our sperm lysate immunoprecipitation data identified Galectin-3 as a candidate glycan-binding protein in this model. Accordingly, we tested the hypothesis that a galectin-3/semenogelin complex contained the HCA epitope by depleting sperm of their surface-bound galectin through competitive inhibition with either lactose, olitigalitin, and asialofetuin. We found that removal of the galectins from the sperm surface reduced HCA-mediated agglutination, and that reduced HCA-mediated agglutination after galectin depletion could be rescued by addition of recombinant Galectin-3 and semenogelin-2 complexes, but not addition of each protein individually. In addition, we found that exogenous addition of recombinant semenogelin to unmodified sperm cultures enhanced HCA-mediated agglutination function. In addition, mass spectrometry-based glycopeptidomic analysis identified several *O*-linked glycosites containing LacNAc on the semenogelin proteins that could serve as binding sites for a galectin. PolyLacNAc sites were not identified as part of the *O*-linked glycosites. Preliminary data indicate that removal of GPI anchored proteins with PI-PLC did not reduce, and in fact enhanced HCA-mediated agglutination, suggesting that the sperm surface *N*-linked glycan that forms the docking site for the HCA-epitope complex is either not a GPI anchored protein, or is a GPI anchor that is resistant to PI-PLC cleavage. GPI anchors modified with an inositol-linked acyl chain (found attached to CD52) have been reported to be resistant to PI-PLC cleavage (*24*). Accordingly, we propose a model of a sperm surface complex consisting of a sperm surface N-glycan, galectin-3, and semenogelin-derived peptides that bind HCA.

Several lines of evidence in the literature support this model: 1) Galectins can anchor highly glycosylated mucins to immune cells and bind extracellular proteins (26, 27). 2) Galectin-1, −3, - 7, and −8 bind human sperm surface glycans, as determined by human lectin array (*28*). 3) Several studies have found Galectin-1 and Galectin-3 on the sperm surface or in seminal plasma by mass spectrometry (*17*, *29–31*). 4) Galectins have been hypothesized to play a role in sperm-egg and sperm-epithelia interactions through an anchoring mechanism that relies on galectin multimerization (*17*, *30*). 5) Reports in the literature indicate that complex *N*-linked glycans on sperm, including those linked to CD52 and CD59, contain polyLacNac moieties that would have a high affinity for galectins (*9*, *32*, *33*). 6) Finally, immunoprecipitation with galectin-3 pulls down semenogelin-1 and −2 from prostasomes (*29*, *31*).

This study has several limitations. Though we isolated a specific tryptic peptide of interest (ISYQSSSTEER) found in seminal plasma and sperm fractions affinity purified on HCA columns, it is not clear which residues of this peptide are critical for binding or whether it was originally part of a longer peptide. Prior studies have reported a CD52-associated semenogelin-1 derived peptide (SQTEKLVAGKSQ)-- that could potentially be continuous with the HCA-purified peptide (*11*). This continuous sequence would be a larger peptide (with the sequence ISYQSSSTEERRLHYGENGVQKDVSQRSIYSQTEKLVAGKSQ) that does carry a LacNAc containing *O*-linked glycosite that could interact with a multivalent galectin-3 on the sperm surface (*34*). However, in this study, we did not determine a specific glycopeptide region of semenogelin that is required for HCA binding. It appears that HCA binding requires a larger environment than simply one glycan, likely a glycopeptide that is presented multivalently as part of a larger complex. Our data indicates that the anchoring of semenogelin to the sperm surface via galectin-3 was required to rescue agglutination after galectin depletion of sperm. However, it still remains unclear whether the HCA epitope involves only semenogelin, or the binding site is a composite target involving portions of both semenogelin and galectin-3. Another limitation is that we did not undertake structural study of the proposed HCA binding site complex. Semenogelin is a largely unstructured and sometimes insoluble protein, and its confirmation is highly reliant on environmental conditions such as the concentration of zinc and other ions, making it difficult to isolate in high concentration and purity for structural studies or to gain any insight on the binding site via computational docking software (*86–88*). Additionally, because semenogelin is physiologically cleaved as part of liquefaction, there is a significant variability in collecting “un-liquefied” samples from human donors. Recombinant semenogelin may address some of these concerns but does not recapitulate the native glycosylation of human derived semenogelin fragments, making it a less feasible alternative to study the structure of HCA-galectin-semenogelin complexes (*18*, *35*). Finally, semenogelin itself is a protein known to multimerize, and this aspect of its biology is not something we explored in this study (*18*). In other contexts, it has been demonstrated that multimerization of glycoproteins can impact galectin binding (*27*). As a result, substantial structural biology remains to be elucidated about the nature of the HCA-semenogelin-galectin complex on the sperm surface. We did not determine the glycoprotein(s) on the sperm surface bound by endogenous Galectin-3 to identify which protein(s) anchor the complex to sperm. In addition, we did not determine if HCA binding is solely reliant on a semenogelin glycopeptide, or whether there is an enhancement of binding when semenogelin is part of a native galectin complex on the surface of sperm. It is possible that a multimeric galectin complex would enable higher affinity avid binding of HCA to multiple semenogelin derived peptides on the sperm surface.

These findings have implications for the mechanism of HCA as a contraceptive and for understanding the biological roles of semenogelin derived peptides and galectins on the sperm surface (*39*, *42*). Semenogelin as a nonhormonal contraceptive target has been proposed in prior literature because single nucleotide polymorphisms in semenogelin have been linked to male factor infertility (*19*, *36*). The role of human semenogelin and semenogelin-derived peptides on sperm function is generally understudied-- there are limited reports using recombinant proteins that suggest a motility inhibition, and/or capacitation modifying function of the intact protein, and there are some studies on semenogelin-derived peptides impacting viral infection in reproductive cells (*18*). Semenogelin derived peptides play a role in sperm motility and anti-adhesion and may even play a role in interaction with cervical mucus, given the ability of HCA to agglutinate sperm and trap them in mucus (*37*). There are limited reports of intracellular semenogelin expression in other cell types, including lung cancers and retina, where the protein may play a different role (*21*, *22*, *38*). However, given the importance of semenogelin glycosylation and physiological cleavage, it is unclear whether studies with recombinant protein recapitulate function in vivo. Given the numerous difficulties in studying the semenogelins, including high levels of glycosylation, low homology of the protein sequence between different species, physiological liquefaction, insolubility, and lack of structural characterization, HCA may be an interesting research tool for biologists interested in this protein or peptides derived from it on the human sperm surface (*90*). In addition, these findings are a first step to finding the exact binding epitope of HCA on sperm and semenogelin, which might lead to improved efforts to affinity-mature the antibody as a second-generation contraceptive product.

## Acknowledgements

We thank Matt Au from Boston University’s Analytical Instrumentation Core and Gregery Woss from Sartorius for technical assistance. We thank the Gummuluru and Muhlberger labs at the Boston University Medical Campus for sharing reagents related to western blots, and Anna Tseng and Aoife O’Connell for technical assistance.

## Author Contributions

SD, DJA, CX, CEC, SF, and SM designed the research. SD, CX, PS, CSW, HMJ, and SF generated data. JP, SD and PS completed statistical analysis of the data. JP and SM obtained IRB approval and provided human samples for the study. SCW, HMJ, and SRS contributed recombinant proteins to the study. KJW produced HCA for the study. CEC, DJA and SM supervised the research and contributed to interpretation of the results. DJA, KJW, CEC, CSW, and SM acquired funding for the study. SD and SF wrote the manuscript. All authors discussed the results and contributed to the final manuscript.

## Ethics declaration

KJW is affiliated with a company that develops antibody-based products, including multipurpose prevention technologies that are contraceptive.

## Data Availability Statement

Data are included in the paper, it’s supplementary files or in public repositories.

## Notes

*Grant support*: NIH P50HD096957 to DJA, NIH T32A1007309 to SD, and K99HL169746 to SCW, S10OD021728 to CEC, National Science Foundation grant No. CHE-2514817 to CEC

### Competing Interest Statement

The authors have declared no competing interest.

