## Supplementary Figures and Tables for "The Human Contraceptive Antibody Targets A Semenogelin Galectin-3 *N-*Linked Glycan Complex on the Sperm Surface"

**Supplementary Figures and Methods**

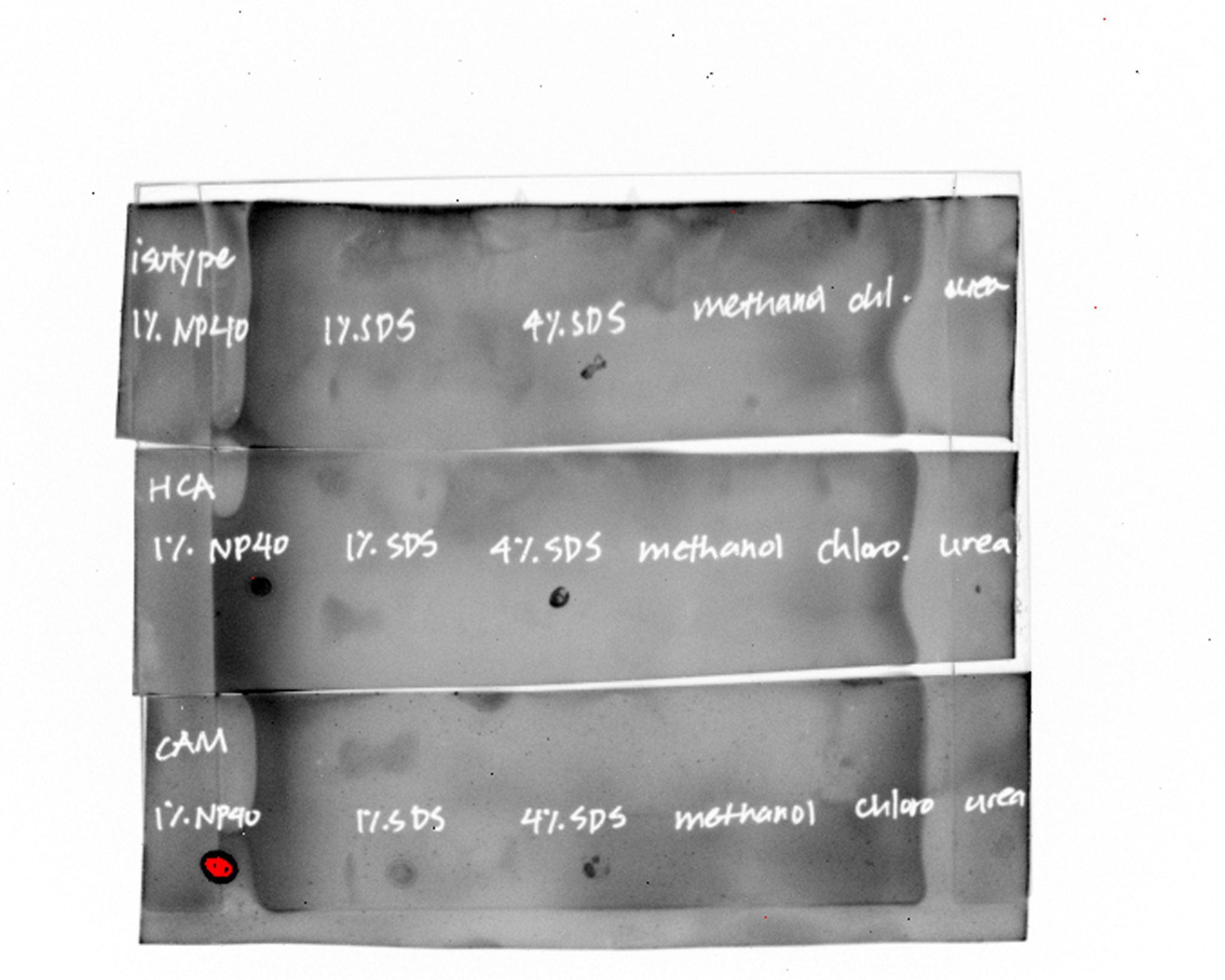

**Supplementary Figure 1: Dot blots of sperm lysates for immunoprecipitation.** Total lysates were blotted by drying 1 uL of lysate onto a PVDF membrane then stained with primary and secondary antibody prior to imaging. Isotope control is VRC01, an anti-HIV antibody.

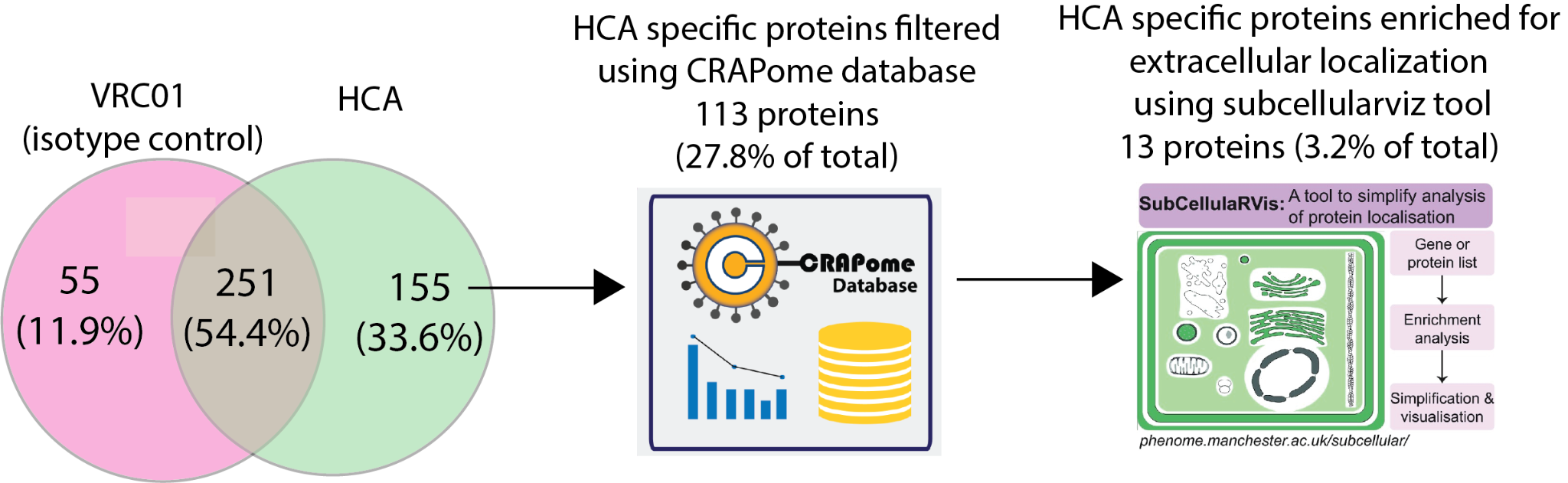

**Supplementary Figure 2: Filtering workflow for sperm lysate immuno-precipitation data.** Proteins identified in HCA affinity-purified sperm lysates after in-gel digest LC-MS/MS were compared to those affinity-purified with an isotype-matched control antibody to filter non-specific interactors. Next, HCA-selected proteins were further filtered by using the CRAPome database to remove common contaminant proteins observed in MS experiments. Proteins identified as a contaminant in over 100 experiments in the database were removed. Finally, the subcellularviz tool was used to enrich extracellularly expressed proteins, given that HCA’s epitope is found on both the sperm surface and secreted into seminal plasma. This led to the list of proteins for which MS data are described in Table 2.

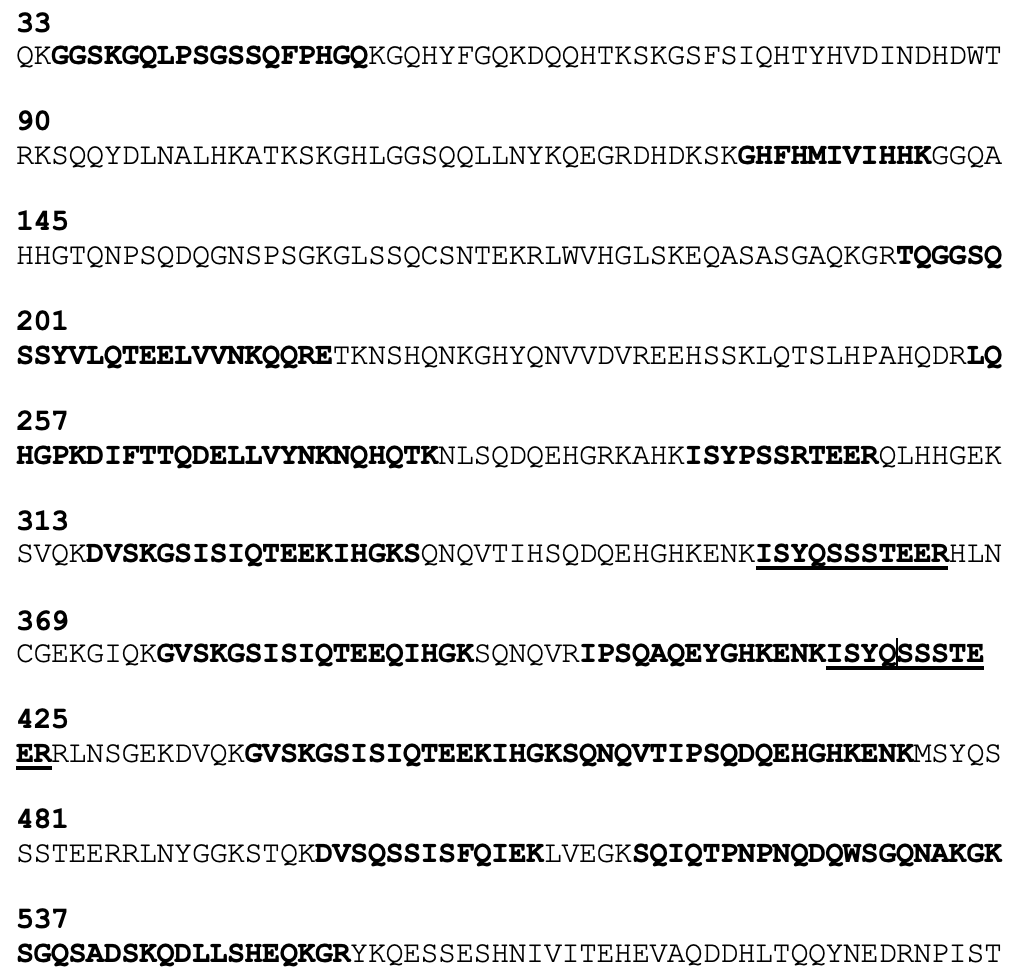

**Supplementary Figure 3: Semenogelin-2-derived peptides marked in the coverage map of peptides identified by mass spectrometry of HCA-purified seminal plasma and sperm lysates.** Peptides identified from seminal plasma HCA immunoprecipitation are bolded, and peptides identified in sperm lysates HCA immunoprecipitation are underlined.

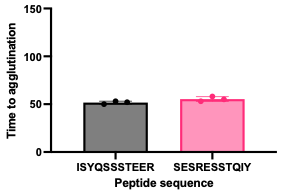

**Supplementary Figure 4: Kinetic Agglutination Competition Assay with ISYQSSSTEER peptide.** 6.25 μg/mL HCA was preincubated with 500 μM peptide in a competition kinetic agglutination assay. ISYQSSSTEER was the peptide isolated from both seminal plasma and sperm affinity purifications after trypsin digestion. SESRESSTQIY is a scramble control peptide containing the same amino acids but in a different sequence.

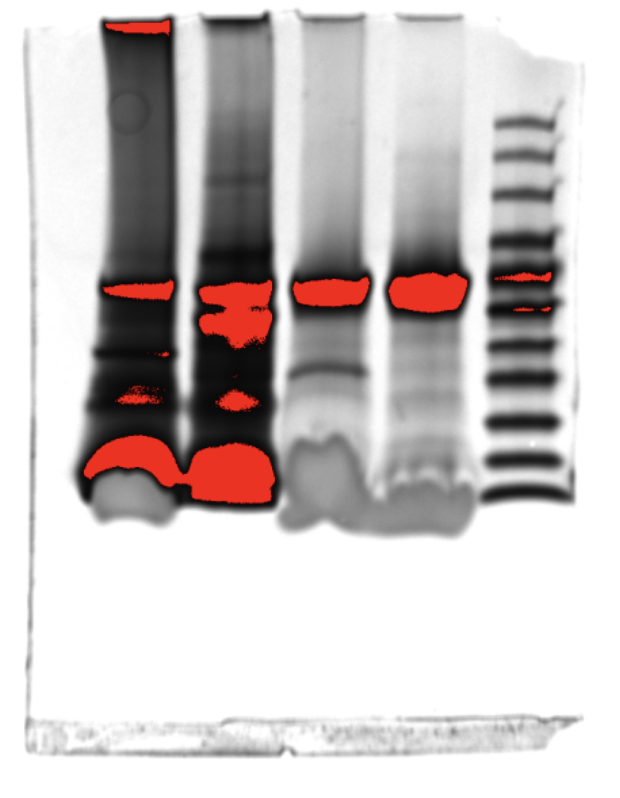

**Supplementary Figure 5: Loading controls for Figure 1E.** SDS PAGE gel of samples in Figure 1 stained with Coomassie. Lanes 1 and 2 are seminal plasma and lanes 3 and 4 are sperm lysate. Lane 5 is a ladder.

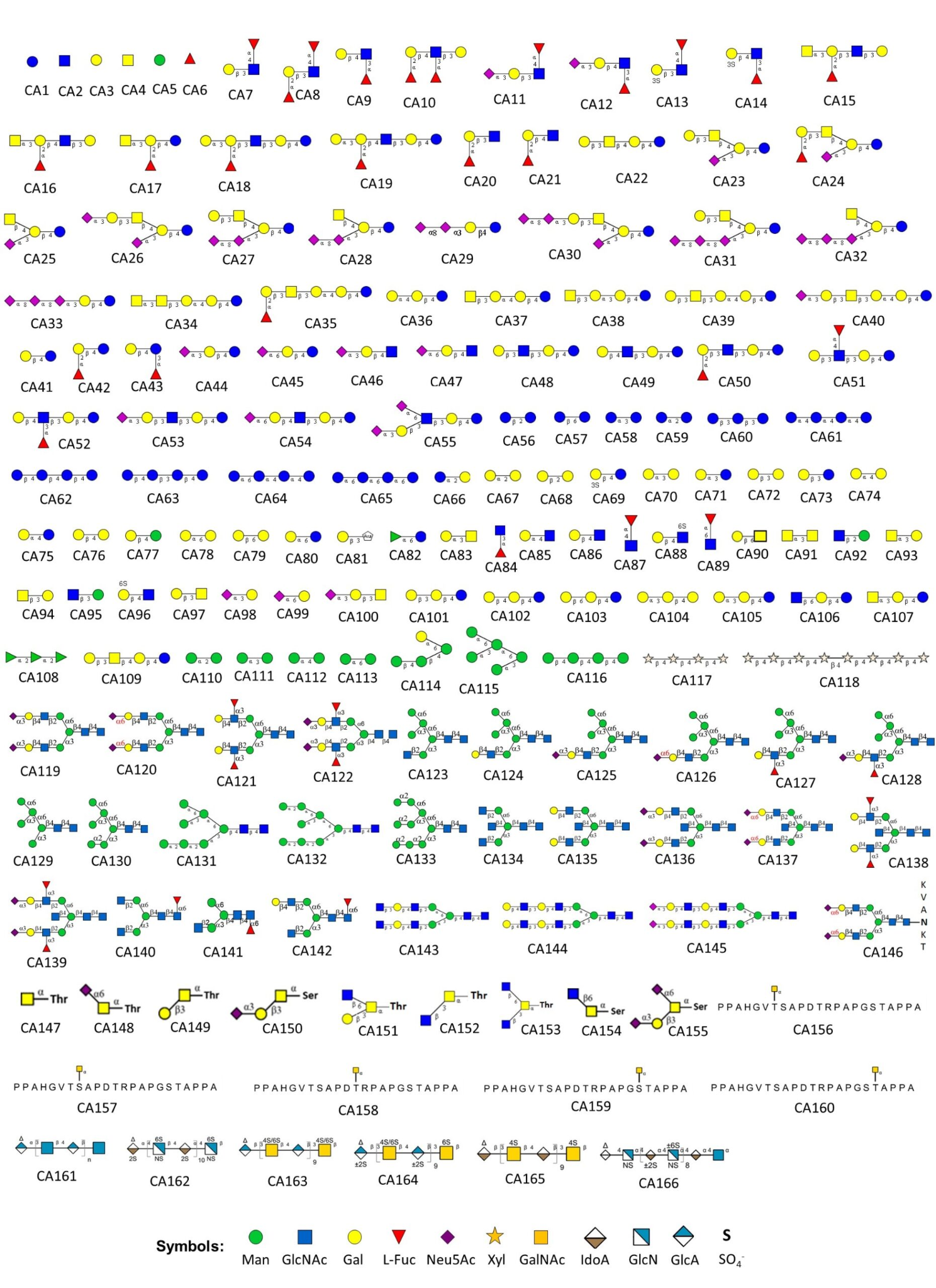

**Supplementary Figure 6: Structures of glycans represented on CatchAll Glycan Array.**

The numbers assigned to each glycan correspond to the x-axis labels in Figure 12. Glycan structures are represented using SNFG (Symbol Nomenclature for Glycans) notation. Adapted from the ZBiotech CatchAll Glycan Array documentation.

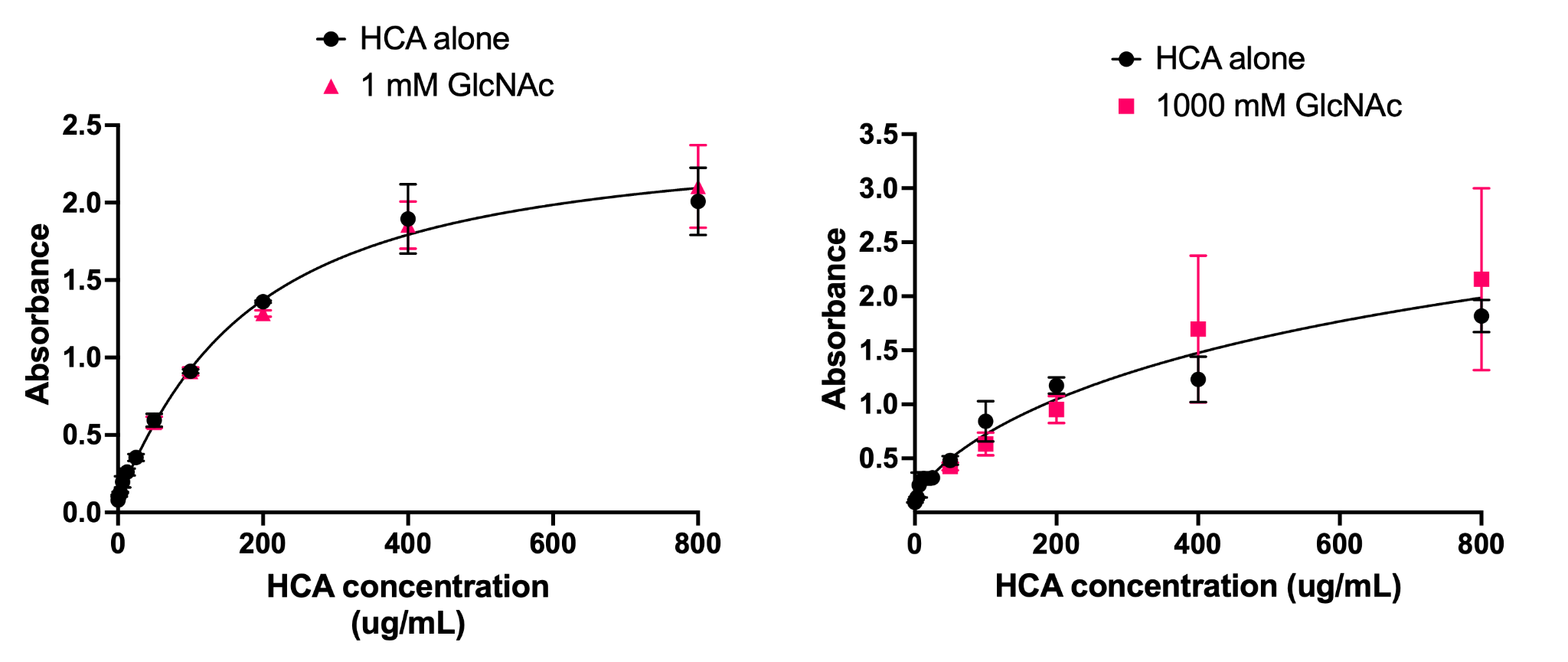

**Supplementary Figure 7: “top hit” on glycan array does not impact HCA binding on whole sperm ELISA.**

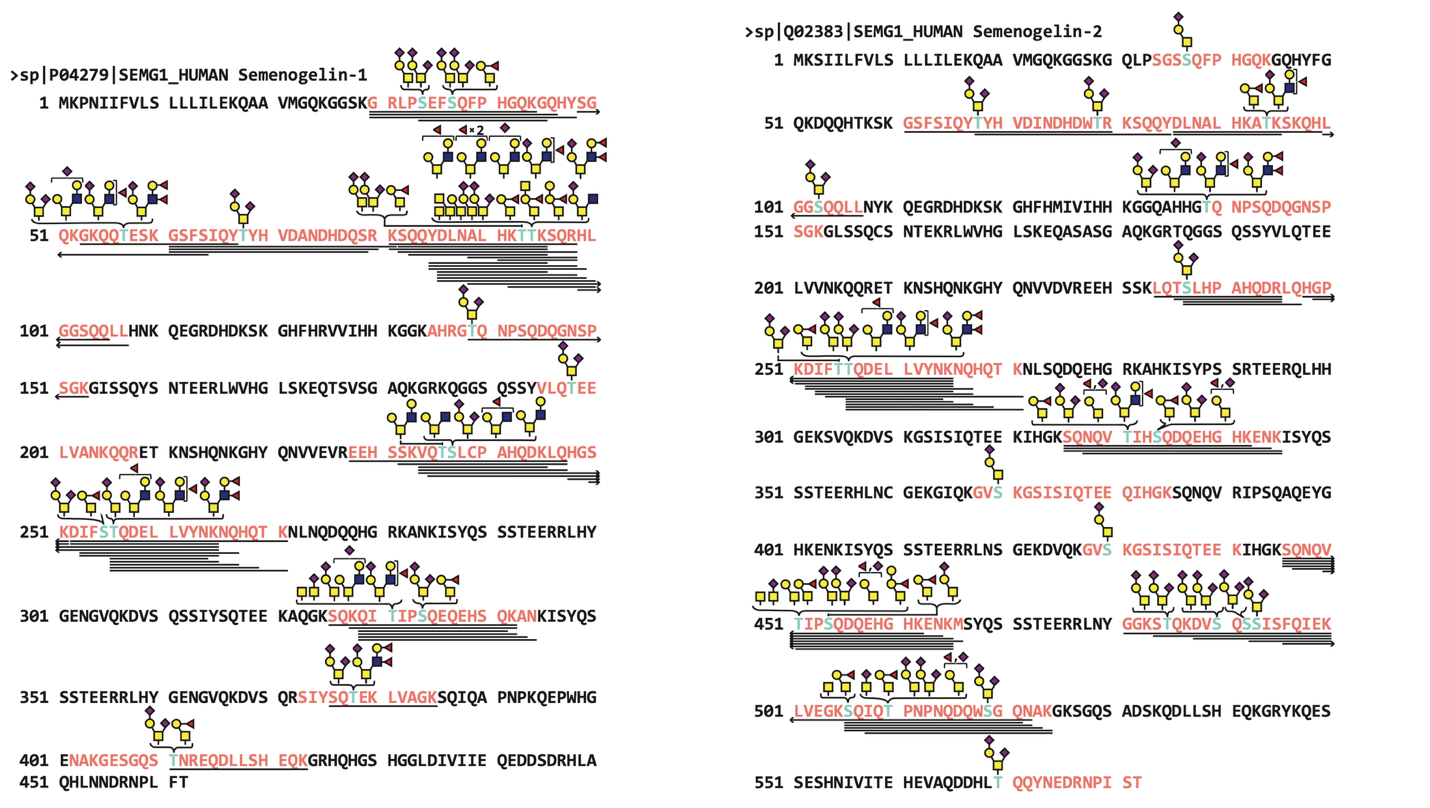
 **Supplementary Figure 8:** GlycoMap of semenogelin-1 and semenogelin-2 as determined by nanoflow liquid chromatography-tandem mass spectrometry.

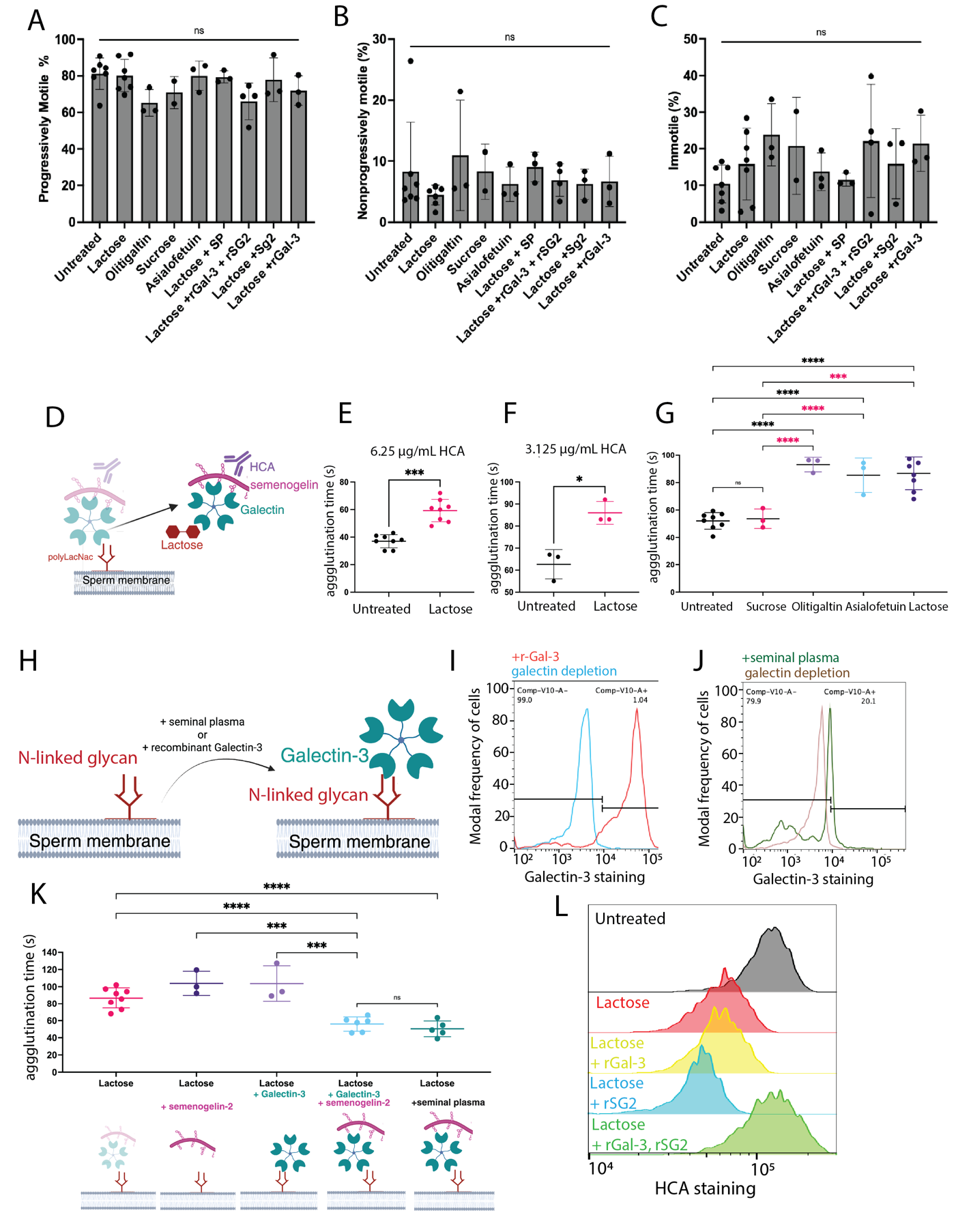

**Supplementary Figure 9:** A) Sperm progressive motility as a percentage after various treatments related to galectin-depletion and rescue. B) Sperm non-progressive motility as a percentage after various treatments related to galectin-depletion and rescue. C) Sperm immotility as a percentage after various treatments related to galectin-depletion and rescue.

**Supplementary Figure 10: HCA-epitope complex schematic.**

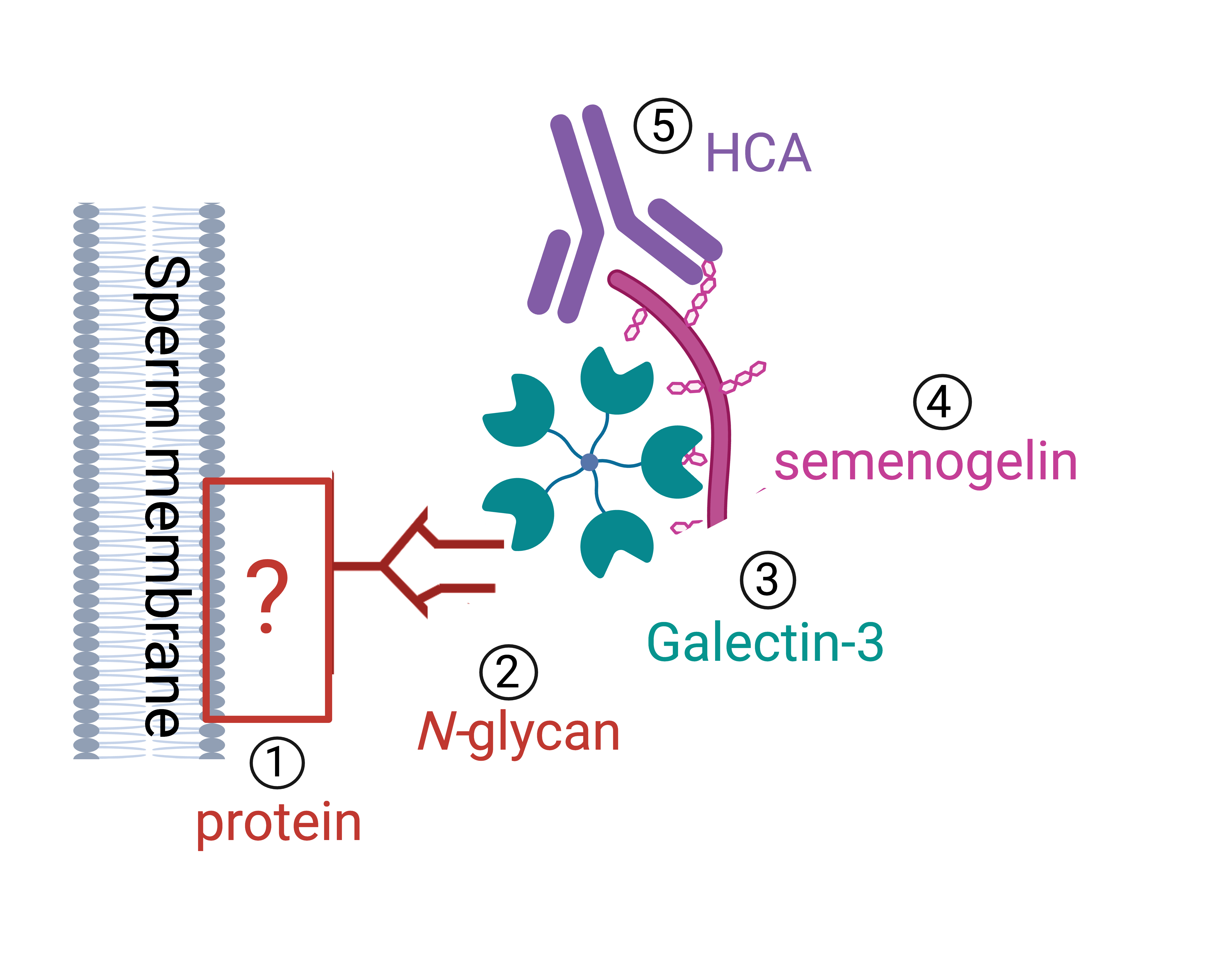

**Supplementary Table 1: Synthesis of data to hypothesize new HCA-epitope model**

| **Model Element** | **Support from literature and this study** | **Link to HCA binding and activity** |
| --- | --- | --- |
| **1: membrane**  **2: N-linked glycans** | Sialidase treatment does not impact HCA binding, so it follows that sialylation of a sperm N-glycan is not required for HCA complex formation and binding **(Purna bioarxiv).** | Removing N-linked glycans from sperm abrogates HCA binding and activity **(Figure 5).**  PI-PLC treatment does not inhibit HCA function and may enhance it. **(Figure 5B)**, Indicating that HCA doesn’t bind a PI-PLC sensitive GPI anchored protein. |
| **3: Galectins** | Galectin -1, -3, -7, -8 bind sperm on array(*1*)  Galectin-1 and -3 found by mass spec on human sperm and in seminal plasma (*2*, *3*)  In our study sperm bind Gal-3 by flow cytometry after exposure to seminal plasma after galectin depletion. | Depleting galectins from the sperm surface reduces HCA-mediated agglutination **(Figure 7).**  Gal-3 was found in sperm lysate IP by HCA **(Table 2).** |
| **4: galectins bind semenogelin** | Gal-3 immuno-precipitation yields semenogelin (*4*)  Semenogelin has many O-linked glycosites that could bind a galectin. (*5*) | Adding recombinant semenogelin restores HCA-mediated agglutination to galectin depletion sperm only in the presence of rGal-3 **(Figure 7).**  Adding recombinant semenogelin enhances HCA mediated agglutination for sperm that are not galectin-depleted **(Figure 6).**  Semenogelin-1 and -2 contain O-linked glycosites with LacNAc moieties that can bind galectins **(Table 3 and 4).** |
| **5: HCA binds semenogelin** | Our western blot data in this study is similar to data from MHS-5, a mouse antibody known to bind human semenogelin (*6*). | HCA IP yields semenogelin **(Table 1 and 2)**  HCA galectin depletion is rescued with addition of both rGal-3 and rSG2 or seminal plasma **(Figure 7).**  HCA binds a large smear that changes sizes on Western blot **(Figure 1).** |

6.	R. J. Evans, J. C. Herr, Immunohistochemical localization of the MHS‐5 antigen in principal cells of human seminal vesicle epithelium. doi: 10.1002/ar.1092140406.

**Supplementary Methods**

Immunopurification of HCA binding partners from seminal plasma: Seminal plasma was centrifuged to pellet debris, and sterile filtered prior to use. Filtered seminal plasma was diluted 1:1 in PBS due to its viscosity and then filtered through a series of spin columns (100KDa, 30KDa) to filter out larger proteins, given that the HCA epitope was presumed to be in the < 20 KDa range. HCA was added to this seminal plasma for a 15-minute pre-incubation in order to pre-form immune complexes, then this mixture was incubated with Protein G spin column for 1 hour with gentle rocking at room temperature. The column was spun in a centrifuge to collect flow through, washed 3x with PBS, then eluants were collected in antibody elution buffer. A gel and western blot was run to confirm purification of the epitope, and then the eluant was spin filtered with 100KDa filter to remove intact antibody. The final eluant was then sent for in-solution digestion and mass spectrometry analysis.

Immunopurification of HCA binding partners from sperm lysates: Motile sperm isolated from several donors was pooled to obtain a total quantity of 1.5 milligrams of total protein. Pellets were resuspended in 1 % NP40, then sonicated on ice for ten minutes using a probe sonicator at the 20% amplitude with a pulse cycle of 10 seconds on and 5 seconds off. This lysate was clarified by centrifugation. 100 µg of antibody was added to each condition, then lysates were incubated at 4 C for immune complex formation overnight. After immune complex formation, samples were incubated with Protein G resin for one hour. The resin was spun down in a centrifuge to remove the non-binding proteins. Resin was washed thrice with PBS, then eluted with SDS. The eluants were separated by SDS PAGE, and gel bands were cut with a sterile razor for mass spectrometry analysis.

Mass spectrometry characterization of affinity purified HCA binding sites from sperm lysates and seminal plasma: Samples were characterized by nano ultra-high-performance liquid chromatography-tandem mass spectrometry (nanoUPLC-MS and MS/MS). Prior to analysis, each sample (1 µg) was reduced with 10 mM of dithiolthreitol, alkylated with 50 mM of iodoacetamide, and then digested with trypsin (C-terminal cleavage at arginine and lysine, 1:50, enzyme: protein) at 37 °C for 16 h and the digest mixture was cleaned up with a C18 SPE cartridge. Next, samples were analyzed by nanoUPLC-MS/MS on an Orbitrap Fusion Lumos Tribrid mass spectrometer (Thermo Fisher Scientific, Waltham, MA, USA) coupled with an ACQUITY UPLC M-Class system (Waters Corp., Milford, MA, USA) via a TriVersa NanoMate (Advion, Ithaca, NY, USA). A nanoEase Symmetry C18 UPLC Trap Column (100 Å, 5 μm, 180 μm × 20 mm, Waters) was used as the trapping column, and a nanoEase MZ HSS C18 T3 UPLC column (100 Å, 1.8 μm, 75 μm × 100 mm, Waters) was used as the analytical column for LC separation. Peptides were trapped at 4 μL/min for 4 min with 1% acetonitrile and 0.1% formic acid (Solvent A). Then, the peptide/glycopeptide mixtures were separated on an analytical column using different concentrations of 99% acetonitrile/0.1% formic acid (Solvent B): 0–1 min: 2% B, 1–3 min: 2–5% B, 3–43 min: 5–40%. MS analyses were performed in the positive mode with the radio frequency (RF) lens set to 30%, and scans were acquired with the following settings: a 120,000 resolution @ *m/z* 200, a scan range of *m*/*z* 370–2000, 1 μscan/MS, a normalized AGC target at 100%, and a maximum injection time of 50 ms. For high energy collisional dissociation (HCD) analyses, initial MS^2^ scans (normalized collision energy (NCE) 27%) were acquired with the following settings: a 15,000 resolution @*m*/*z* 200, a scan range of *m/z* 100–2000, 1 μscan/MS, AGC target 1 × 10^6^, and a maximum injection time of 100 ms. MS/MS data were searched against 20,352 entries in a UniProtKB database restricted to Homo sapiens (downloaded in May 2021) by PMI-Byonic (version v3.8-11, Protein Metrics Inc., Cupertino, CA, USA). Carbamidomethylation (Cys) was set as a fixed modification, whereas Met oxidation was defined as variable modifications. For glycosylation analysis, the Protein Metrics 132 human *N-*glycan and 6 most common *O-*glycan library were used for the assignment of *N*- and *O-* linked glycosylation, respectively. The mass tolerance was set to 10 and 20 ppm at the MS and MS/MS levels, respectively. Enzyme specificity was set to C-terminal of arginine and lysine, with a maximum of two missed cleavages.

Mass spectrometry of seminal plasma O-glycopeptidome: 10 kDa and 30 kDa Ultracentrifugal molecular weight cut-off (MWCO) filters (Vivacon® 500, Hydrosart) were first rinsed three times with 500 µL of 20 mM tris pH 7.4 by centrifuging the filters at 6,000 g for 5 min. Filtered seminal plasma containing approximately 1 mg of protein content, as determined by Nanodrop, was thawed and immediately loaded onto the filter. The samples were incubated at 37 °C for 1 hr. After incubation, the seminal plasma was diluted with 200 µL of 20 mM tris, pH 7.4, and filtered through either a 30 kDa or a 10 kDa filter by centrifugation at 10,000 g for 10 minutes. The retentate was diluted with 200 µL of tris and centrifuged once more. To the eluate that was filtered at 30 kDa, DTT was added to a concentration of 2 mM and the mixture was incubated at 65 °C for 1 h, followed by alkylation with 5 mM IAA for 15 min in the dark at RT. 1 unit of O-Glycoprotease IMPa (NEB, P0761S) was then added to the 30 kDa sample and incubated at 37 °C for 3 hrs, followed by addition of 200ng of trypsin (Promega, V5111) and incubated for another 3 hrs. After digestion (30 kDa) or filtration (10 kDa), the samples were acidified by adding 4 μL of formic acid and desalted using 10 mg Strata-X 33 μm polymeric reversed-phase SPE columns (Phenomenex). The SPE columns were activated using 1 mL of acetonitrile followed by 1 mL of 0.1% formic acid, 1 mL of 0.1% formic acid in 40% acetonitrile, and equilibrated with 1 mL of 0.1% formic acid. After equilibration, the samples were added to the column and rinsed twice with 200 μL of 0.1% formic acid. The columns were transferred to a 1.5 mL tube for elution using two 150 μL additions of 0.1% formic acid in 40% acetonitrile. The eluent was then dried using a vacuum concentrator.

Samples were analyzed by online nanoflow liquid chromatography-tandem mass spectrometry using an Orbitrap Eclipse Tribrid mass spectrometer (Thermo Fisher Scientific) coupled to a Dionex UltiMate 3000 HPLC instrument (Thermo Fisher Scientific). Samples were resuspended in 10 µL of 0.1% formic acid (solvent A), and 6 µL were injected onto an Acclaim PepMap 100 column packed with 2 cm of 5 μm C18 material (Thermo Fisher, 164564). Peptides were then separated on a 15 cm PepMap RSLC EASY-Spray C18 column packed with 2 μm C18 material (Thermo Fisher, ES904) using a linear gradient from 0 to 35% solvent B (0.1% formic acid with 80% ACN) in 60 min with a constant flow rate of 300 nL/min. Full scan MS1 spectra were acquired at a resolution of 60,000 full width at half maximum (FWHM) at 400 m/z, an automatic gain control (AGC) target of 3e5, and a mass range from 300 to 1,500 m/z. Dynamic exclusion settings allowed 2 repeats in 7 s before exclusion for 7 s. Only charge states 2 to 6 were selected for fragmentation. MS2s were generated with a cycle time of 3 s. Fragmentation scans were collected in the Orbitrap at 7,500 resolution. Higher-energy collisional dissociation (HCD) was performed on all selected precursor masses with the following parameters: isolation window of 2 m/z, stepped collision energy 25–30–40% normalized collision energy (nCE), Orbitrap detection (resolution of 7,500), maximum inject time of 50 ms, and a standard AGC target. An additional electron transfer dissociation (ETD) fragmentation of the same precursor was triggered if 1) 3 of 8 HexNAc or NeuAc fingerprint ions (126.055, 138.055, 144.07, 168.065, 186.076, 204.086, 274.092, and 292.103) were present at ± 0.1 m/z and greater than 5% relative intensity, and 2) the precursors were present at a sufficient abundance for localization using ETD. If the precursor was between 300–850 m/z and over 3e4 intensity, an ETD scan was triggered and read out using the ion trap at a normal scan rate. For precursors between 850–1500 m/z with an intensity greater than 1e5, Supplementary activation was used (EThcD) was also applied at 15% nCE and read out in the Orbitrap at 7500 resolution. Both used charge-calibrated ETD reaction times, 150 ms maximum injection time, and 200% standard injection targets.

RAW files were first searched using Byonic v4.0.12.66 (ProteinMetrics), against the human proteome (UP000005640) without modifications or a glycan database to create a focused protein database. Due to the wide specificity of endogenous proteases, the searches were performed with fully non-specific cleavage and any number of missed cleavages. The mass tolerance was set to 10 and 20 ppm at the MS and MS/MS levels, respectively. The outputs of the initial search were filtered for peptides with a score above 150 and a logprob above 1.5, and a focused protein database was generated from the peptide identifications above the cutoff. The second search was performed with the following settings: RAW files were searched against the focused protein database, with the 22-common O-glycan library set as a common modification, allowing 1. O-glycans that also appear in the 9 most common O-glycan library were set to an allowance of 2. The total number of common modifications was set to 2. The output of the final search was filtered for peptides having a glycan, a score above 150, and a logprob above 1.5. All reported glycopeptides above the cutoffs were manually validated.

Supplementary Data Files 1: Mass spectrometry outputs from sperm lysates IP as excel files for HCA

Supplementary Data Files 2: Mass spectrometry outputs from sperm lysates IP as excel files for CAMPATH

Supplementary Data Files 3: Mass spectrometry outputs from sperm lysates IP as excel files for VRC01

Supplementary Data Files 4: Mass spectrometry outputs from unpurified sperm lysates in-gel digest as excel files HCA

Supplementary Data Files 5: Mass spectrometry outputs from unpurified sperm lysates in-gel digest as excel files CAMPATH
